# High-throughput transcriptomics and benchmark concentration modeling for potency ranking of per- and polyfluoroalkyl substances (PFAS) in exposed human liver cell spheroids

**DOI:** 10.1101/2020.10.20.347328

**Authors:** A.J.F. Reardon, A. Rowan-Carroll, S.S. Ferguson, K. Leingartner, R. Gagne, B. Kuo, A. Williams, L. Lorusso, J.A. Bourdon-Lacombe, R. Carrier, I. Moffat, C.L. Yauk, E. Atlas

**Affiliations:** Environmental Health Science and Research Bureau, Healthy Environments and Consumer Safety Branch, Health Canada; Biomolecular Screening Branch, Division of the National Toxicology Program, National Institute of Environmental Health Sciences National Institutes of Health; Chemicals and Environmental Health Management Bureau, Healthy Environments and Consumer Safety Branch, Health Canada; Water and Air Quality Bureau, Healthy Environments and Consumer Safety Branch, Health Canada; Department of Biology, University of Ottawa, Ottawa, ON, Canada; Department of Biochemistry, University of Ottawa, ON, Canada

**Keywords:** High-throughput transcriptomics, new approach methodologies, benchmark concentration, perfluoroalkyl substances, biological potency

## Abstract

Per- and polyfluoroalkyl substances (PFAS) are some of the most prominent organic contaminants in human blood. Although the toxicological implications from human exposure to perfluorooctane sulfonate (PFOS) and perfluorooctanoate (PFOA) are well established, data on lesser-understood PFAS are limited. New approach methodologies (NAMs) that apply bioinformatic tools to high-throughput data are being increasingly considered to inform risk assessment for data-poor chemicals. The aim of this investigation was to identify biological response potencies (i.e., benchmark concentrations: BMCs) following PFAS exposures to inform read-across for risk assessment of data-poor PFAS. Gene expression changes were measured in primary human liver cell microtissues (i.e., 3D spheroids) after 1-day and 10-day exposures to increasing concentrations of 23 PFAS. The cells were treated with four subgroups of PFAS: carboxylates (PFCAs), sulfonates (PFSAs), fluorotelomers, and sulfonamides. An established pipeline to identify differentially expressed genes and transcriptomic BMCs was applied. We found that both PFCAs and PFSAs exhibited a trend toward increased transcriptional changes with carbon chain-length. Specifically, longer-chain compounds (7 to 10 carbons) were more likely to induce changes in gene expression, and have lower transcriptional BMCs. The combined high-throughput transcriptomic and bioinformatic analyses supports the capability of NAMs to efficiently assess the effects of PFAS in liver microtissues. The data enable potency ranking of PFAS for human liver cell spheroid cytotoxicity and transcriptional changes, and assessment of *in vitro* transcriptomic points of departure. These data improve our understanding of the health effects of PFAS and will be used to inform read-across for human health risk assessment.

## Introduction

Per and polyfluoroalkyl substances (PFAS) are synthetic fluorinated contaminants that have a variety of commercial and industrial applications. A long history of manufacture in parallel with thermal and chemical stability contribute to the widespread distribution of PFAS detection of perfluoroalkyl acids (PFAAs) in humans (Hansen *et al.*, 2001) and wildlife (Giesy and Kannan, 2001). Common sources of exposure to the broader class of PFAS include diet (Fromme *et al.*, 2007; Bjermo *et al.*, 2013; Vestergren *et al.*, 2012), indoor air and dust (De Silva *et al.*, 2012; Makey *et al.*, 2017; Shoeib *et al.*, 2011), and contact with soils at PFAS-contaminated sites (e.g., aqueous film-forming foams (AFFFs)) (Hu *et al.*, 2016). To date, guidelines have been proposed to regulate perfluorooctane sulfonate (PFOS) and perfluorooctanoate (PFOA), the two most prominent PFAS in Canadian drinking water (Health Canada (HC), 2018a, 2018b). However, there exists a lack of data to make regulatory recommendations for other, lesser-known PFAS and their precursors.

Regulatory practices for human health risk assessment currently rely on resource-intensive animal data. However, there is a push among international regulatory agencies to harness new approach methodologies (NAMs) to rapidly acquire information on data poor chemicals such as PFAS while reducing reliance on animal models (Kavlock *et al.*, 2018). *In vitro* toxicological data has been used previously as a prediction of potency and observed effects by applying toxicokinetic models to calculate an administered equivalent dose (AED) using *in vivo* to *in vitro* extrapolation for *in vivo* potency ranking (Pham *et al.*, 2019). Although AEDs can be determined for the most prominent PFAS (e.g., PFOS and PFOA) toxicokinetic data are not available for most PFAS. However, other approaches such as benchmark concentration (BMC) modeling (Crump, 1984; Slob *et al.*, 2005) may be used to identify thresholds of molecular change (Paul Friedman *et al.*, 2020). In this approach, a BMC, defined as a change in response from a predetermined threshold (e.g., 1 standard deviation) relative to the background incidence, was determined from continuous concentration-response data Combined with high-throughput transcriptomics, BMC models can be advantageous in providing quantitative toxicogenomic information for various risk assessment applications when adequate toxicokinetic data is not available.

In a companion paper, toxicological assessments of four prototype PFAS with TempO-Seq profiling were examined in primary human liver cell spheroids (Rowan-Carroll *et al.* submitted). Transcriptomic profiles were used to develop an analytical pipeline for application toward investigating a larger group of PFAS. Within this previous work, exposure to PFOS, PFOA, short chain perfluorobutane sulfonate (PFBS), and long-chain perfluorodecane sulfonate (PFDS) resulted in transcriptional responses and pathway perturbations. A bioinformatics pipeline demonstrated the capability of high-throughput gene expression analyses to establish transcriptomic patterns for read-across application. Common upstream regulators were identified (e.g., peroxisome proliferator-activated receptor PPARα) and pathway perturbations, including fatty acid oxidation, cholesterol biosynthesis, and stearate biosynthesis were revealed. These observations were in agreement with past *in vivo* (Bijland *et al.*, 2011; Curran *et al.*, 2008; Das *et al.*, 2017; Menger *et al.*, 2020; Wan *et al.*, 2012) and *in vitro* (Bjork *et al.*, 2011; Hickey *et al.*, 2009; Naile *et al.*, 2012) studies revealing PFAS-induced changes in expression of specific genes relating to important processes in the liver.

The overarching goal of this investigation was to identify biological response potencies (i.e., BMCs) following PFAS exposures to inform read-across in order to inform risk assessment of data-poor PFAS present at federal contaminated sites where AFFFs have been used, impacting surrounding soils and groundwater. Here, we expand on our previous work from Rowan-Carroll et al. (submitted) building on the generated data and developed bioinformatics approach to evaluate the potency of 19 additional data-poor PFAS with BMC modeling. For comparison, PFAS were subdivided into perfluoroalkyl carboxylates (PFCAs), perfluoroalkyl sulfonates (PFSAs), fluorotelomers, and sulfonamides (Table 1) and included the data for PFBS, PFOS, PFDS, and PFOA from the initial study (Rowan-Carroll *et al.* submitted). Biological activity was evaluated by determining the number of differentially expressed genes (DEGs) and transcriptional BMCs for each PFAS. Previously, liver-toxicity thresholds based on gene expression profiles from chemical exposure has been established in HepaRG cells (Ramaiahgari *et al.*, 2019), and these data were used to create an *in vitro* potency rank-order for PFAS in our liver spheroids based on transcriptional concentration-response effects. The potency rank-order of PFAS were compared using the overall median BMCs of transcripts as well as the lowest 5^th^ percentile gene BMCs, and an *in vitro* point of departure (POD) was proposed for each PFAS.

**Table 1.**
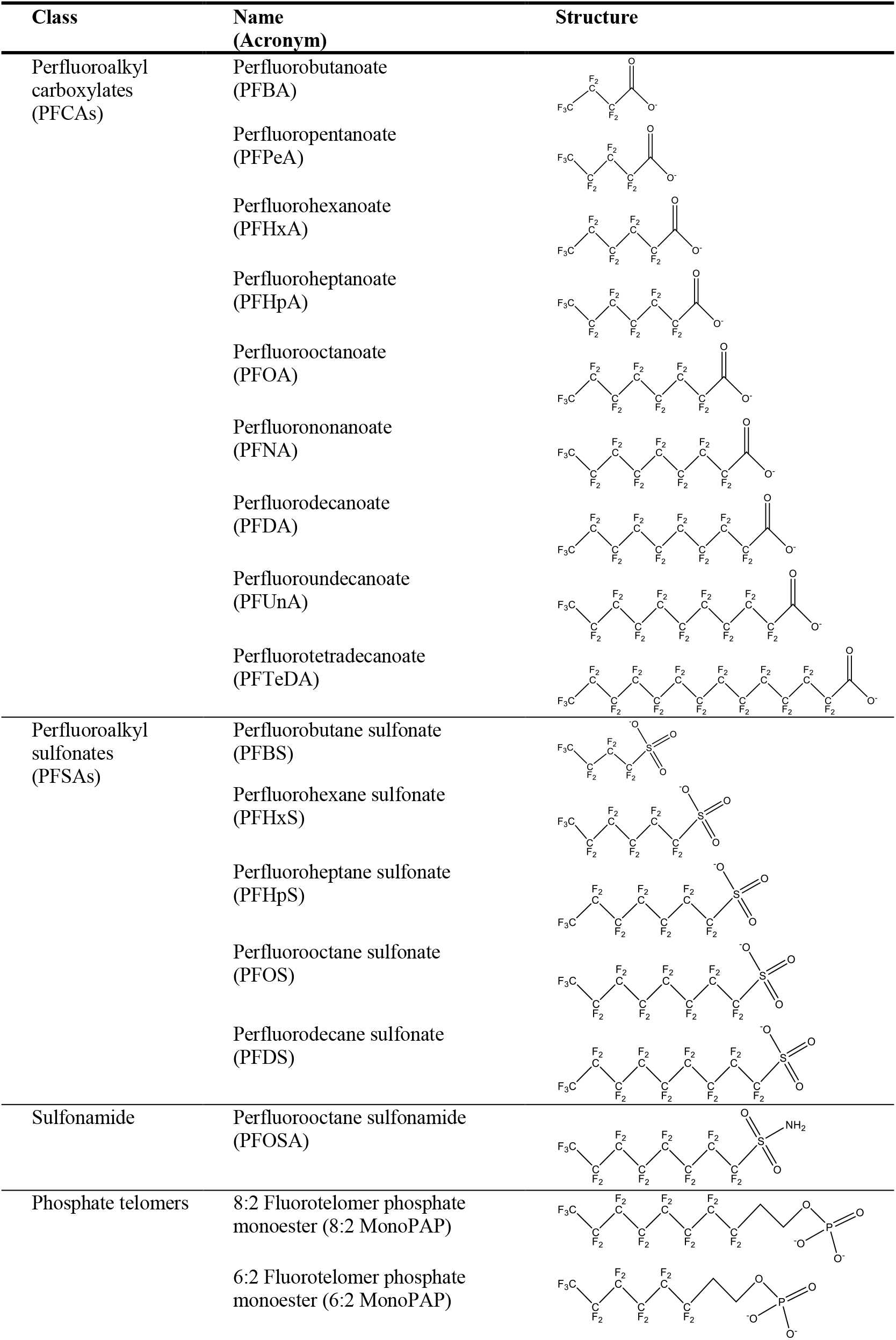
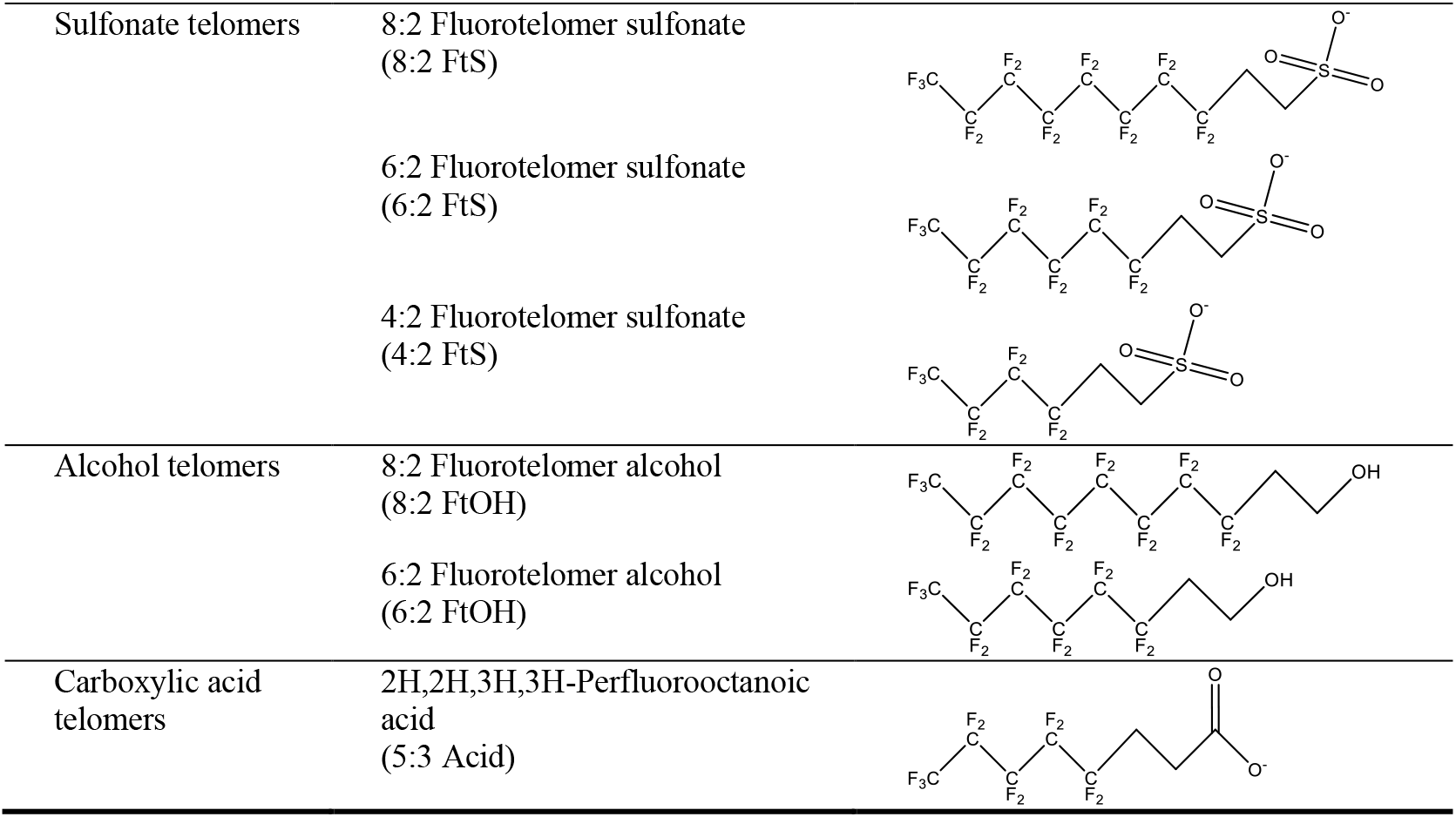
List of names and formulae, and structures of project-specific PFAS

| Class | Name<br>(Acronym) | Structure |
| --- | --- | --- |
| Perfluoroalkyl<br>carboxylates<br>(PFCAs) | Perfluorobutanoate<br>(PFBA) |  |
|  | Perfluoropentanoate<br>(PFPeA) |  |
|  | Perfluorohexanoate<br>(PFHxA) |  |
|  | Perfluoroheptanoate<br>(PFHpA) |  |
|  | Perfluorooctanoate<br>(PFOA) |  |
|  | Perfluorononanoate<br>(PFNA) |  |
|  | Perfluorodecanoate<br>(PFDA) |  |
|  | Perfluoroundecanoate<br>(PFUnA) |  |
|  | Perfluorotetradecanoate<br>(PFTeDA) |  |
| Perfluoroalkyl<br>sulfonates<br>(PFSAs) | Perfluorobutane sulfonate<br>(PFBS) |  |
|  | Perfluorohexane sulfonate<br>(PFHxS) |  |
|  | Perfluoroheptane sulfonate<br>(PFHpS) |  |
|  | Perfluorooctane sulfonate<br>(PFOS) |  |
|  | Perfluorodecane sulfonate<br>(PFDS) |  |
| Sulfonamide | Perfluorooctane sulfonamide<br>(PFOSA) |  |
| Phosphate telomers | 8:2 Fluorotelomer phosphate<br>monoester (8:2 MonoPAP) |  |
|  | 6:2 Fluorotelomer phosphate<br>monoester (6:2 MonoPAP) |  |
| Sulfonate telomers | 8:2 Fluorotelomer sulfonate<br>(8:2 FtS) |  |
|  | 6:2 Fluorotelomer sulfonate<br>(6:2 FtS) |  |
|  | 4:2 Fluorotelomer sulfonate<br>(4:2 FtS) |  |
| Alcohol telomers | 8:2 Fluorotelomer alcohol<br>(8:2 FtOH) |  |
|  | 6:2 Fluorotelomer alcohol<br>(6:2 FtOH) |  |
| Carboxylic acid telomers | 2H,2H,3H,3H-Perfluorooctanoic acid<br>(5:3 Acid) |  |

## Methods

### Cell culture

3D InSight™ Human Liver Microtissues were purchased from InSphero (Brunswick, ME) in a 96 well format, with a single spheroid per well. These spheroids are a co-culture model from 10 different human liver donors and are a metabolically active system of hepatocytes and Kupffer cells. Upon arrival, culture media were replaced with InSphero human liver maintenance medium–tox, and spheroids were acclimated for 24 hours prior to PFAS exposures.

### Chemical preparation and exposure conditions

Scoping PFAS (PFBS, PFOS, PFDS, PFOA) were the subject of a previous investigation and the details of their purchase and preparation are described there (Rowan-Carroll *et al.* submitted). The remainder of PFAS chemicals were donated by the US Environmental Protection Agency. A complete list of chemical structure and formula information are presented in Table 1. PFAS were dissolved in tissue grade dimethyl sulfoxide (DMSO, Sigma-Aldrich, Oakville, ON). Working stock solutions were prepared in quadruplicate and ranged in concentration from 0.02 to 100 mM stocks and diluted 1:1000 in culture media. For a few higher concentrations of PFAS with low solubility, the dilution factor was 1:300, and thus, the DMSO controls were at 0.3%. The concentration of DMSO was determined based on the solubility of individual PFAS (DMSO in exposure medium ranged from 0.1% to 0.3%) but was kept constant across the exposure range for each PFAS. Under circumstances that required highest DMSO content to dissolve PFAS in solution (e.g., above 0.1 %) DMSO concentration equivalent controls were concurrently run to account for relative matrix-induced effects.

Concentration selection for PFAS was based on several factors. The US EPA’s ToxCast program uses a top concentration of 100 μM for *in vitro* assays and this concentration was anticipated to yield a robust genomic response required for BMC models. Thus, this was selected as our top concentration. A large range of concentrations were selected up to 100 μM to ensure that robust transcriptomic responses were achieved across all PFAS that minimized potential for cytotoxic concentrations, and maximized potential to span each gene-level BMC. Final concentrations for all PFAS in this study were: 0.2, 2, 10, 20, 50, 100 μM unless otherwise stated. Concentrations were selected in order to capture elicited effects and responses in liver microtissues from a broad range of PFAS, and the range of exposures aligned with previous high-throughput screening studies within the US EPA ToxCast program (Judson *et al.*, 2010).

Primary human liver cell spheroids were cultured in 3D InSight™ Human Liver Microtissue Media (InSphero, Brunswick, ME). Exposure media were created by addition of individual PFAS at the indicated concentrations for 10 days at 37 °C and 5% CO_2_ (N = 4). Refreshed media containing PFAS were replaced every 3 days and spent media were kept frozen until subsequent cytotoxicity analysis. Prior to lysis at day-1 or day-10, liver spheroids were washed with Dulbecco's phosphate-buffered saline (DPBS) (Thermo Fisher Scientific, Franklin, MA) and then lysed using 5-7 μl of TempO-Seq lysis buffer. Liver microtissues were triturated and left to lyse for approximately 10 min, then frozen at −80 °C. For the 10-day exposure, spent media was replaced every 3 days with freshly prepared media containing the various concentrations of PFAS.

### Spheroid cytotoxicity

Cytotoxicity was determined using the lactate dehydrogenase (LDH) assay (LDH-Glo, PROMEGA J2380, Madison, WI), as per manufacturer’s instructions. Briefly, at the indicated times after the initiation of treatment, supernatants were diluted 1:10 in storage buffer (200 mM Tris-HCl pH 7.3 containing 10% glycerol and 1% bovine serum albumin) and kept frozen at −80 °C. At the time of analysis samples were diluted with an equal volume of LDH Detection Reagent and equilibrated for 60 minutes at room temperature before reading the luminescence (GloMax 96 Microplate Luminometer, Promega Corp, Madison, WI) in relative luminescence units (RLU). Treatment effects on cytotoxicity were determined after performing statistical analysis of the RLU data after blank subtraction using the Kruskal-Wallis one-way analysis of variance followed by the Dunn’s test for multiple comparisons with Sigma Plot 13™ for both Day 1 and Day 10 (N = 4). The sample RLU was converted to a ratio by dividing by the average RLU of the DMSO controls and calculating the mean and standard deviations of these ratios for each treatment and time point. A list of PFAS exposures that were removed from induced cytotoxicity is presented in the supplementary information (Table S1).

### TempO-Seq library building and next generation sequencing

Gene expression was measured using the human TempO-Seq S1500+ panel (Mav *et al.*, 2018) (BioSpyder Technologies Inc, Carlsbad, CA). This panel is comprised of 3000 genes selected by the NIEHS to represent a biologically diverse set of pathways that are environmentally responsive (https://federalregister.gov/a/2015-08529).

For TempO-Seq analysis, liver spheroids treated with increasing concentrations of PFAS or vehicle control DMSO over 24 hours or 10 days were lysed *in situ* using a volume of 2x TempO-Seq lysis buffer equal to the residual volume of media. Lysates and positive controls (Human Universal Reference RNA - uhrRNA (Agilent Cat# 740000), and Human Brain Total RNA brRNA (ThermoFisher AM7962)), as well as no-cell negative controls (1X TempO-Seq lysis buffer alone) were hybridized to the detector oligo mix following the manufacturer’s instructions (TEMPO-SEQ Human Tox +Surrogate with Standard Attenuation Transcriptome Kit (96 Samples) BioSpyder Technologies, Inc. Carlsbad, CA, USA). Hybridization was followed by nuclease digestion of excess oligos, detector oligo ligation, and amplification of the product with the tagged primers according to manufacturer’s instructions. At the time of amplification, each sample was ligated to a sample-specific barcode that allows for combining samples for sequencing purposes. Labelled amplicons were then pooled and purified using NucleoSpin Gel and PCR Clean-up kits, (Takara Bio USA, Inc, Mountain View, CA). Libraries were sequenced in-house using a NextSeq 500 High-Throughput Sequencing System (Illumina, San Diego, CA) using 50 cycles from a 75-cycle high throughput flow cell.

### Data processing and identification of Differentially Expressed Genes (DEGs)

TempO-Seq data were extracted from the bcl files and demultiplexed (i.e., assigned to respective sample files) with bcl2fastq v. 2.20.0.42. The fastq files were processed with the “pete. star. script_v3.0” supplied by BioSpyder. Briefly, the script uses STAR v.2.5 to align the reads and the qCount function from QuasR (Gaidatzis *et al.*, 2015) to extract the feature counts specified in a gtf file from the aligned reads.

The data were then passed through internal quantity control scripts. Boxplots of the log2 CPM (counts per million) were plotted to ensure a reproducible distribution between replicates within a group. Read counts were also used to identify overtly cytotoxic concentrations alongside results from the LDH assay. Samples with low read counts were often associated with high levels of cytotoxicity (LDH release). Hierarchical clustering plots were generated (hclust function: default linkage function of hclust function in R; complete-linkage) for all the samples per time point using a distance metric defined as 1-Spearman correlation in order to identify potential outliers. Low count probes (i.e., probes that did not have a median of 5 counts in at least one group) were flagged as absent probes and eliminated from the data set. To account for differences in the percentage of DMSO used, the data were normalized to matched controls and rescaled to the average of the DMSO 0.1% dose group.

DEG analysis was conducted on the counts using the default parameters of DESeq2 v1.24 with respective control and exposure groups. Probes reaching the threshold of a false discovery rate (FDR) adjusted p-value ≤ 0.05 and an absolute fold change ≥ 1.5 (on a linear scale) were flagged (probes passing filters; PPF) and were retained for further analyses.

Heatmaps were generated with the pheatmapfunction (Kolde, 2019) from R with the row and column distances defined as 1-Spearman correlation respectively. Values were averaged between replicates; CPM mean values of PPFs were averaged between replicates within each group. Resulting values were then log2 transformed. Principal components were calculated with the prcompR function. The percentages of variation on the axes were gathered from the “importance” field produced by the summary function applied to the resulting object of the prcomp function. The work above was done with R v.3.6.1. Further details on pipeline analysis and QA/QC are described in the supplementary information. PCA plots were generated with averaged PPF values.

### Benchmark concentration analysis

BMC modeling was conducted using the BMDExpress (v.2.3) software package that is freely available on-line (https://github.com/auerbachs/BMDExpress-2/releases). The BMDExpress platform rapidly analyzes large datasets for determining models of best-fit. Additional information on BMDExpress and details of BMC analysis is available at https://github.com/auerbachs/BMDExpress-2/wiki/Benchmark-Dose-Analysis (Yang *et al.*, 2007).

Imported data underwent pre-filtering using the William’s Trend Test with 100 permutations and a 1.5-foldchange (FC) cut-off to select probes (i.e., genes) that exhibit significant concentration-response behaviour (p < 0.05). Data exhibiting an increasing or decreasing change in response were modeled using listed best-fit models, including Hill, Power, Linear, Polynomial (2° and 3°), and Exponential (2, 3, 4, 5) to identify potential concentration-response relationships and best-fit BMC. For linear and polynomial models, the Nested Chi-square test was used to determine best-fit models followed by a comparison of Akaike information criterion (AIC) for nested models (Hill and Power) with a goodness of fit p-value of 0.1. Hill models were flagged if the ‘k’ parameter was 1/3 the lowest positive concentration, and in the case of a flagged Hill model, the next best-fit model with a goodness of fit p-value > 0.05 was selected. Additional parameters included a benchmark response factor of 1 SD relative to the response of controls, a confidence level of 0.95, power restricted to ≥ 1, and maximum of 250 iterations.

To avoid potential bias that may result from the inclusion of poor-fit genes, the following probe filters were included: a goodness of fit p-value < 0.1, BMCU/BMCL ≥ 40, as well as data producing BMC values > the highest non-cytotoxic concentration (i.e., 100 μM). BMC gene accumulation plots reflecting the distribution of gene BMCs across the range of concentrations, and gene BMC estimates from best-fit models were exported from BMDExpress. A robust central measure of the transcriptional activity induced by a chemical was defined as the median BMC of all filtered genes adhering to best-fit models and an in-house bootstrap R-script producing 95 % confidence intervals (median BMC ± 95 % CI).

### BMC gene accumulation plots and established thresholds of liver injury

BMC accumulation plots have been used for hazard identification of potentially liver-toxic compounds. Here, we use the approach previously described by Ramaiahgari *et al.*, 2019 (detailed in the supplementary information). Given that the Ramaiaghari *et al.* thresholds were derived from 2D HepaRG cells and a training set of non-liver toxic chemicals, we could not use these same thresholds for identifying liver toxicity herein. Therefore, the published approach from Ramaiahgari *et al.* was used to rank order potentially liver-toxic PFAS and compare potencies. Gene accumulation plots after PFAS exposure of human liver cell spheroids were generated using the parameters and filters from Ramaiahgari *et al.* to qualitatively identify potentially liver-toxic chemicals (# genes with BMCs ≥ 105) and again to rank order the toxicity based on transcriptional responses.

### BMC models to derive transcriptomic points of departure

We sought to identify a transcriptomic POD that was not based on a pre-determined set of genes (e.g., a pathway database). For the purposes of this investigation the terms ‘gene’ and ‘transcript’ were used synonymously. The lower bound 5th percentile of all gene BMCs that was rounded to the gene with an integer value closest to the 5th percentile estimate was used for analysis. The gene BMC, upper (BMCU), and lower (BMCL) bound values were identified from exported best models from BMDExpress. A similar methodology (5th percentile) has been used in previous work to represent a lower bound estimate within the ToxCast database typically to derive values for the bioactivity exposure ratio (BER) (Friedman *et al.*, 2019). Each gene was checked to ensure that none of the BMC values included were fit with a “flagged” Hill model for which no other best-fit model with a p-value < 0.05 could be obtained.

## Results

### Cytotoxicity assessment of PFAS in liver cell spheroids

The enzyme lactate dehydrogenase (LDH) normally present in the cytoplasm of liver cells is extensively released into culture media when the cell membranes are disrupted. Thus, detection of LDH leakage into culture media is an established marker of cellular cytotoxicity. We found no evidence of cytotoxicity for short-chain PFAS (C4 – C6) in the liver cell spheroids after 10 days of exposure up to the 100 μM concentrations investigated (Fig. 1). For PFCAs, 1-day exposure to the top concentration of PFDA (100 μM) and the highest two concentrations of PFUnA (50 and 100 μM) elicited increases in LDH levels. Ten-day exposures to longer-chain (C9–C14) PFCAs – PFNA, PFDA and PFUnA increased LDH release from mid to high concentrations (Fig. 1A). Although most PFSAs and fluorotelomers were non-cytotoxic, PFOS and sulfonamide PFOSA caused cytotoxicity at concentrations above 20 μM (Figs. 1B and 1C respectively). Cytotoxic responses from PFOS and PFOSA exposure were not influenced by time as the same concentrations from 1-day exposure were also cytotoxic after 10 days for both PFAS. Within our internal quality controls coinciding with LDH assays, the number of mapped reads (mapped to S1500+ probe set) was also considered for identifying overtly cytotoxic samples that were subsequently removed from further analyses.

**Figure 1.**
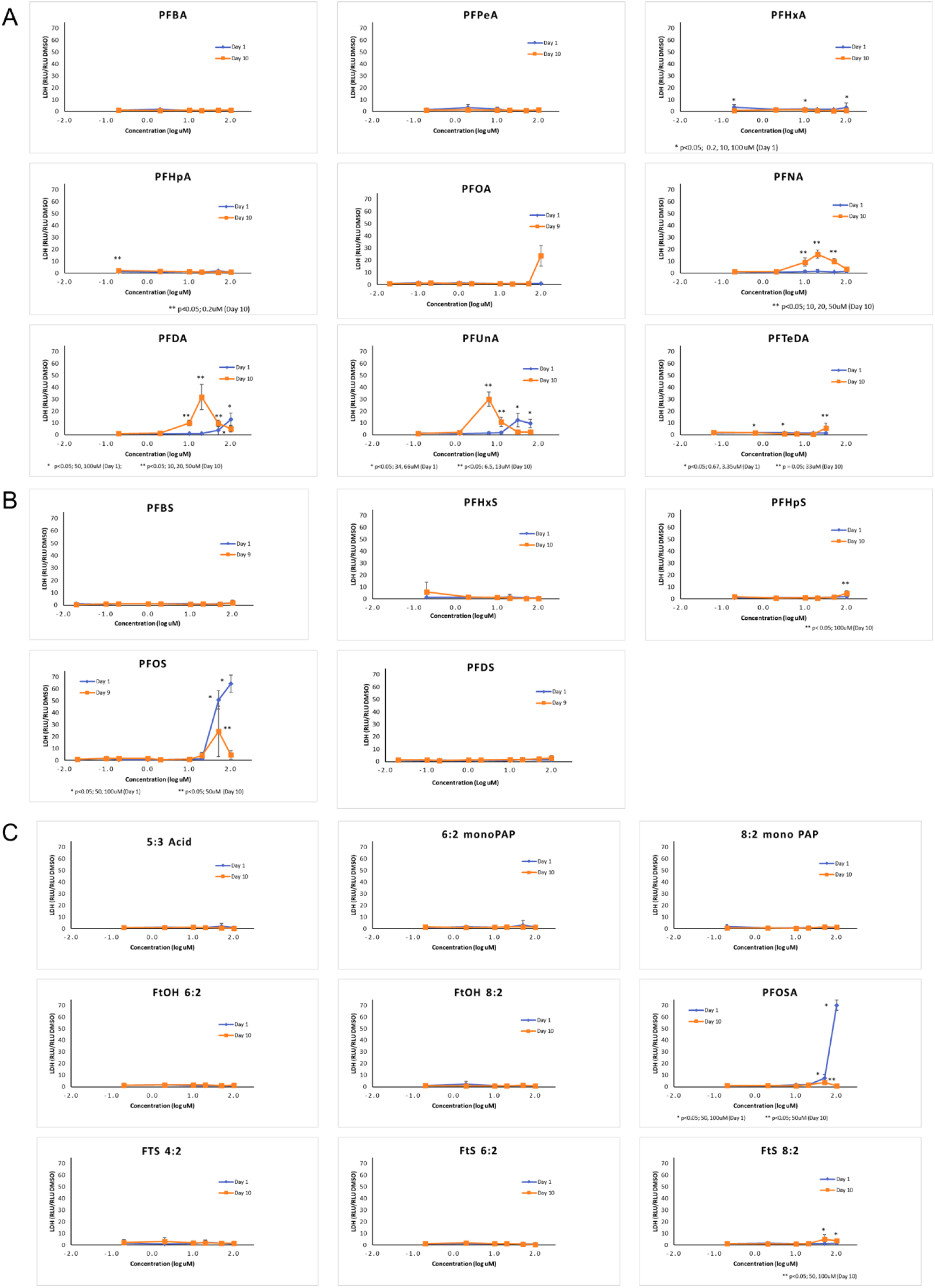
Cytotoxicity analysis (LDH release) of liver spheroids following exposure to increasing concentrations of PFAS: A) carboxylates, B) sulfonates, and C) fluorotelomers/sulfonamides from 1-day (blue circles) and 10-day (orange squares) exposure. Different from controls, p < 0.05 (1-Day) * Different from controls, p < 0.05 (10-Day) **

### Transcriptomic data QA/QC – overview, cytotoxicity and outliers

A quality assessment was completed prior to analysis of differentially expressed genes as described in Rowan-Carroll *et al.* (submitted) with water-only controls and reference standards within TempO-Seq plates. In brief, observations were conducted confirming low (< 0.3 %) signal observed within blank control wells without cells/RNA, and positive controls of commercial human universal reference RNA (two per plate) with high Spearman correlation coefficients (> 0.94) across all plates. The median and minimum mapped reads within the study were 2,000,000, and 400,000 reads, respectively and inter-run variability as a result of plate-effects were corrected for (example calculation in supplementary information).

Observations of cytotoxicity from LDH assays were paired with read counts from TempO-Seq analyses to further identify cytotoxic concentrations to be removed from subsequent transcriptomic analyses. Of 1392 total samples, cytotoxicity was identified within 124 samples from relatively high exposure concentrations of PFDA, PFUnA, PFOS, and PFOSA at 1-day, and from PFOA, PFNA, PFDA, PFUnA, PFTDeA, PFOS, PFOSA, and 8:2 FtS at 10-day exposures and removed (concentrations listed in Table S1). An additional 103 samples had either a low number of mapped reads or were identified as outliers from hierarchical clustering for a total of 227 treatment samples removed from subsequent analyses. Consolidating all these approaches led to sample sizes of n=3 or n=4 for each PFAS experimental treatment group.

### Identifying differentially expressed genes (DEGs)

DEGs were defined as transcripts with |FC| ≥ 1.5, and FDR p ≤ 0.05 relative to time-matched solvent controls (Table 2). A trend-line of log-transformed number of DEGs was established to show the increasing number of DEGs with PFAS exposure and is depicted in the supplementary information (Table S2). Most short-chain PFCAs (C4-C6) and the longest-chain PFCA (PFTeDA) had no discernable trend from either exposure time, except for PFBA at 10 days having an increased number of DEGs at higher (50 and 100 μM) exposures. In contrast, spheroids exposed to long-chain PFCAs, from 7-carbon PFHpA to 11-carbon PFUnA, exhibited increasing numbers of DEGs with concentration. It was not possible to discern trends for PFNA, PFDA, and PFUnA at the 10-Day time point because of cytotoxicity at relative low exposure concentration (> 2 μM) as higher exposure concentrations led to shut down of cell transcription following overt cytotoxicity for some PFAS. PFSAs followed the same trend as PFCAs: long-chain PFSAs had increasing numbers of DEGs with increasing concentration at both time points. PFOSA exposure caused a concentration-dependent increase in DEGs until inducing cytotoxicity at 50 μM; however, most PFAS fluorotelomers had no discernable trend, either yielding no DEGs or low numbers of DEGs observed at all concentrations.

**Table 2.** Differentially expressed genes and concentration-responses of human liver spheroids after 1-day and 10-day exposure to PFAS

|  | Day | Concentration (μM) |  |  |  |  |  |
| --- | --- | --- | --- | --- | --- | --- | --- |
|  |  | 0.2 | 2 | 10 | 20 | 50 | 100 |
| PFBA | 1 | 6 | 65 | 1 | 26 | 15 | 58 |
|  | 10 | 41 | 49 | 17 | 19 | 132 | 199 |
| PFPeA | 1 | 52 | 1 | 0 | 2 | 0 | 0 |
|  | 10 | 34 | 40 | 22 | 40 | 26 | 24 |
| PFHxA | 1 | 119 | 43 | 22 | 60 | 101 | 24 |
|  | 10 | 123 | 167 | 139 | 61 | 44 | 62 |
| PFHpA | 1 | 16 | 0 | 3 | 1 | 15 | 72 |
|  | 10 | 55 | 10 | 19 | 25 | 57 | 91 |
| PFOA | 1 | 0 | 11 | 72 | 71 | 229 | 491 |
|  | 10 | 2 | 11 | 85 | 106 | 625 | -* |
| PFNA | 1 | 37 | 2 | 40 | 182 | 239 | 786 |
|  | 10 | 165 | 36 | -* | -* | -* | -* |
| PFDA | 1 | 1 | 4 | 0 | 62 | 353 | -* |
|  | 10 | 29 | 34 | -* | -* | -* | -* |
| PFUnA<sup>A</sup> | 1 | 40 | 20 | 119 | 227 | -* | -* |
|  | 10 | 0 | 0 | -* | -* | -* | -* |
| PFTeDA<sup>B</sup> | 1 | 46 | 9 | 128 | 55 | 69 | 43 |
|  | 10 | 42 | 0 | 67 | 47 | 67 | -* |
| PFBS | 1 | 0 | 34 | 7 | 0 | 50 | 72 |
|  | 10 | 0 | 2 | 0 | 7 | 55 | 79 |
| PFHxS | 1 | 129 | 0 | 0 | 10 | 12 | 18 |
|  | 10 | 0 | 1 | 2 | 8 | 22 | 60 |
| PFHpS | 1 | 12 | 2 | 17 | 27 | 63 | 230 |
|  | 10 | 80 | 10 | 93 | 132 | 262 | 467 |
| PFOS | 1 | 1 | 50 | 171 | 295 | -* | -* |
|  | 10 | 4 | 64 | 175 | 498 | -* | -* |
| PFDS | 1 | 0 | 17 | 49 | 75 | 177 | 190 |
|  | 10 | 0 | 42 | 140 | 181 | 241 | 237 |
| 5:3 Acid | 1 | 0 | 3 | 0 | 12 | 10 | 20 |
|  | 10 | 15 | 3 | 6 | 6 | 20 | 131 |
| 6:2 MonoPAP | 1 | 0 | 0 | 28 | 1 | 7 | 11 |
|  | 10 | 5 | 17 | 15 | 27 | 30 | 20 |
| 8:2 MonoPAP | 1 | 1 | 0 | 0 | 0 | 2 | 0 |
|  | 10 | 6 | 0 | 0 | 0 | 0 | 11 |
| 6:2 FtOH | 1 | 12 | 55 | 43 | 21 | 67 | 32 |
|  | 10 | 93 | 41 | 107 | 22 | 10 | 0 |
| 8:2 FtOH | 1 | 4 | 11 | 0 | 13 | 0 | 0 |
|  | 10 | 37 | 20 | 13 | 31 | 20 | 72 |
| 4:2 FTS | 1 | 4 | 6 | 21 | 14 | 3 | 6 |
|  | 10 | 47 | 40 | 71 | 31 | 33 | 43 |
| 6:2 FtS | 1 | 3 | 5 | 15 | 6 | 69 | 7 |
|  | 10 | 34 | 40 | 12 | 10 | 16 | 51 |
| 8:2 FtS | 1 | 4 | 29 | 56 | 127 | 212 | 237 |
|  | 10 | 81 | 88 | 136 | 172 | 234 | -* |
| PFOSA | 1 | 20 | 53 | 22 | 148 | -* | -* |
|  | 10 | 38 | 102 | 93 | 488 | -* | -* |

\* Overtly cytotoxic concentrations eliminated from analysis, <sup>A</sup> Concentration range of PFUnA (0.13, 1.3, 6.5, 13, 34, 66 μM), <sup>B</sup> Concentration range of PFTeDA (0.06, 0.67, 3.35, 6.7, 17, 33 μM)

There were similar trends of increasing DEGs with exposure concentration over both observed time-points. However, longer exposure time (10 days) generally led to a higher number of DEGs compared to shorter exposures (e.g., 20 μM PFOS elicited 295 and 498 DEGs over 1 and 10 days, respectively). Moreover, cytotoxicity was more prominent with longer exposure durations (Fig. 1), indicating that specific PFCAs (PFOA, PFNA, PFDA, PFUnA, and PFTeDA), as well as PFOS, PFOSA and 8:2 FtS would likely elicit increased transcriptional activity (i.e., affecting a greater number of genes) with increased exposure duration if not for overt cytotoxicity at 10-days (Table 2). For example, exposure to PFNA adversely affect the spheroids at relatively low concentrations (> 2 μM). The increased number of DEGs with increased exposure is anticipated to be consistent with accumulation and alteration of a greater number of biological pathways. Although this is beyond the scope of the current study, investigation into the corresponding biological pathways will be the subject of future investigation. Finally, there were instances where the peak number of DEGs occurred at low to moderate concentrations rather than at the highest non-cytotoxic concentration: PFPeA at 0.2 μM (1-day), PFHxA at 0.2 μM (1-day), PFHxS at 0.2 μM (1-day), PFTeDA at 10 μM (1-day), and 6:2 MonoPAP at 20 μM (1-day).

### Benchmark concentration modelling

#### Overall trends in concentration-response

Concentration response curves from high-throughput transcriptomic data were produced using the transcriptomic workflow described by Philips *et al.* for BMDExpress v2.3 (Phillips *et al.*, 2019). Each gene BMC represents the concentration where transcript levels deviated from the baseline of background concentration by the predesignated benchmark response (1 standard deviation). The median transcriptomic BMC of all genes that could be modeled within the considered S1500+ probe set was then calculated from the responses of multiple genes each with an individual lower (BMCL) and upper (BMCU) limit. Confidence intervals (CI) were determined from resampling using a bootstrap R-script to create 95 % CI for the entire set of genes with BMCs. Forest plots of the median gene BMC (± 95 % CIs) for the total subset of genes where a BMC estimate was modeled from the overall S1500+ gene-set for each PFAS exposure are depicted in Figure 2. A full list of all PFAS BMCs (median ± 95 % CI) over each exposure time is detailed in the supplementary information (Table S3).

**Figure 2.**
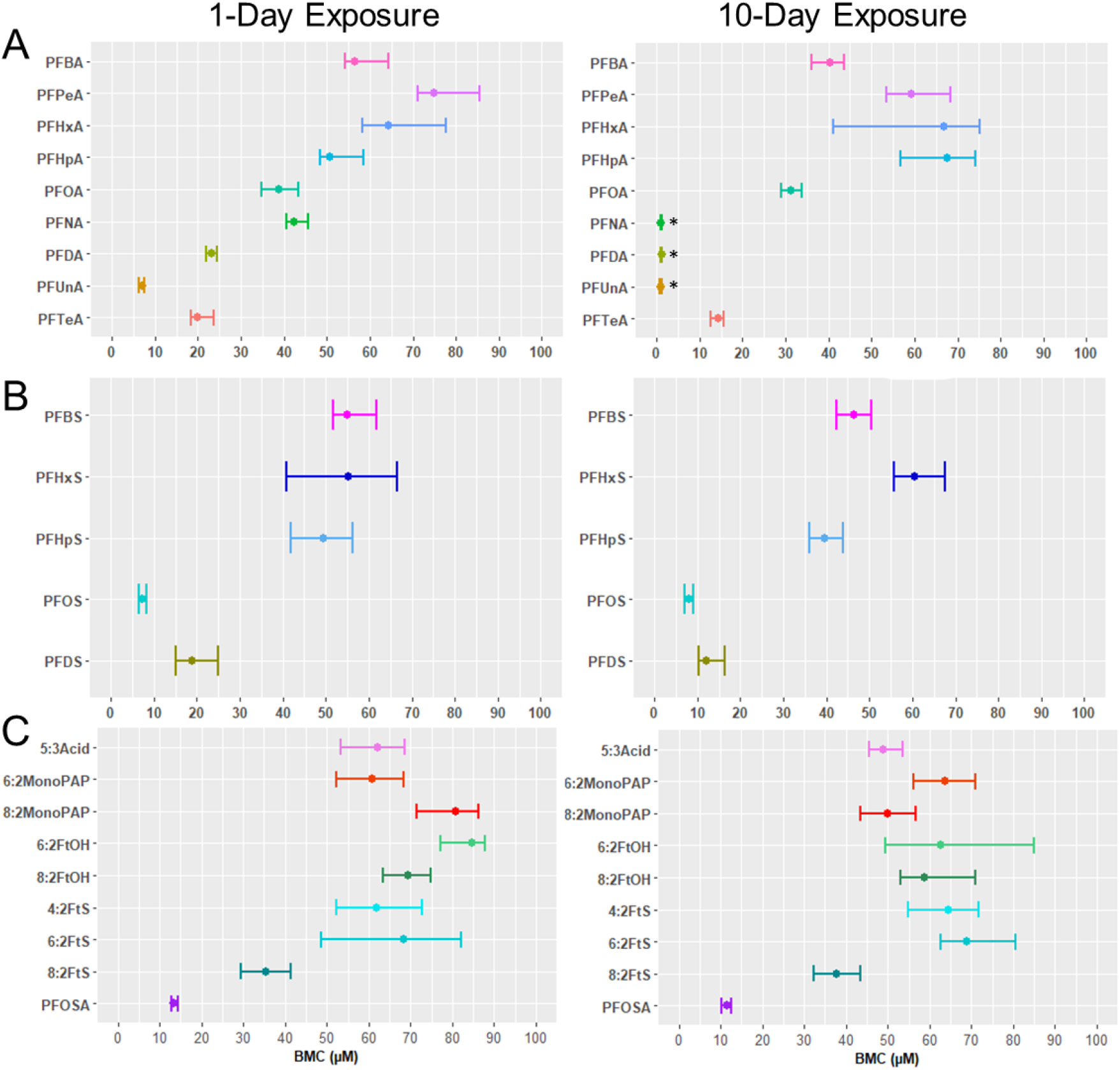
Benchmark concentration (BMC) responses (median +/− 95 % CI) of human liver spheroids exposed to A) PFCAs, B) PFSAs, or C) fluorotelomers/sulfonamides exposed for 1-day and 10-days. The analysis was conducted in BMDExpress 2.3. *BMCs derived from two concentrations only due to overt cytotoxicity at low exposure concentrations

Potency for the PFCAs and PFSAs (based on the median gene BMC) corresponded to PFAS structure; specifically, the data suggest decreasing BMC values (increasing potency) with increasing carbon chain length (Fig. 2). PFAS from 4 to 7 carbons were generally less potent than longer-chain PFAS, with PFUnA being the most potent (1-day BMC 7.0 ± 1.3 μM), followed by PFOS (1-day BMC 7.3 ± 0.8 μM), PFDS (10-day BMC 12.0 ± 6.1 μM), and PFTeDA (10-day BMC 14.3 ± 3.2 μM). Of fluorotelomers and sulfonamides, the most potent was the PFOS-precursor PFOSA (10-day BMC 11.5 ± 2.4 μM). The trend of PFAS potency with chain length was similar between the two time points. However, it was noted that BMC modeling could only be applied to the lowest two concentrations at 10-days for PFNA (0.2 and 2 μM), PFDA (0.2 and 2 μM), and PFUnA (0.13 and 1.3 μM) due to overt cytotoxicity.

#### Gene accumulation from concentration-response models

Gene-level accumulation plots show the individual genes with BMCs creating a profile of biological effects in the liver cell spheroids from PFAS exposure (Fig. 3). Within accumulation plots, the gene order is plotted sequentially on the y-axis with each gene’s BMC on the x-axis. The most sensitive genes have the lowest BMC values, and a steep curve represents increased transcriptional activity over a small concentration range. Alternative log-transformed accumulation plots are provided for interpretation within the supplementary information (Figs. S1 and S2). Some curves are shorter than others because overtly cytotoxic concentrations were not included in gene accumulation plots. Overall, for PFCAs and PFSAs, the trend of biological activity followed the same pattern as the overall median gene BMC above, with steeper slopes to the left of the plots with increasing chain-length, suggesting increasing activity/potency with aliphatic chain length.

**Figure 3.**
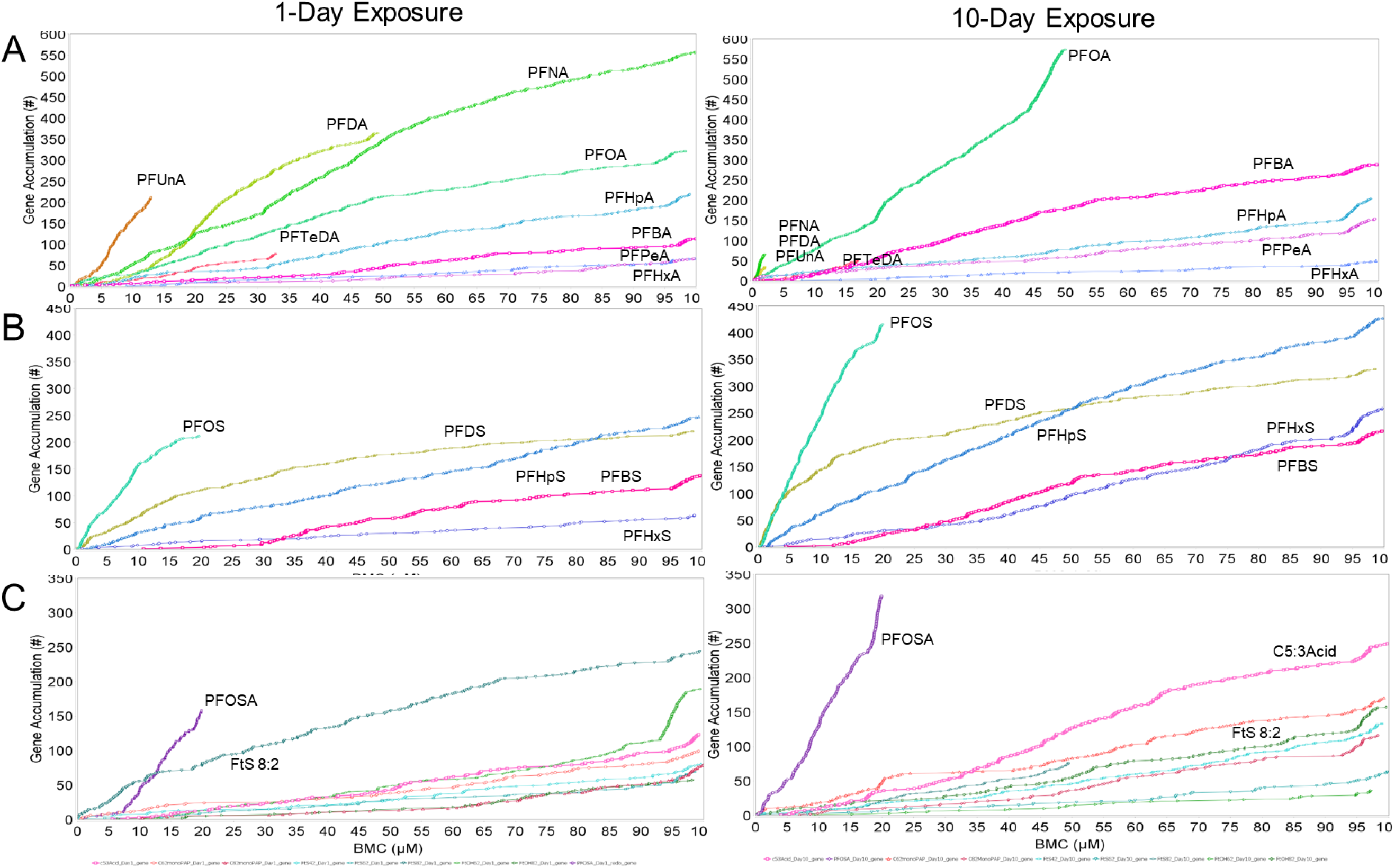
Gene accumulation plots of BMC responses in human liver spheroids following exposure to A) PFCAs, B) PFSAs, or C) fluorotelomers/sulfonamides exposed from 1-day and 10-days. The analysis was conducted in BMDExpress 2.3.

On day 1, decreased transcriptional activity (lower number of accumulated gene BMCs) was observed for short-chain PFAS (PFBA, PFPeA, and PFHxA) than longer-chain PFCAs (PFHpA, PFOA, and PFNA). Note that the 10-carbon (PFDA) and 11-carbon (PFUnA) curves are cut off at lower BMCs because of cytotoxicity (Fig. 3); however, at any given non-cytotoxic concentration, these longer-chain PFAS exhibit more biological activity than the short chain. By 10 days, the trend of increasing gene BMC accumulation is not as clear due to increased cytotoxicity, particularly from exposure to PFNA, PFDA, and PFUnA (Fig. 3). Some PFAS elicited increased transcriptional activity after 10-days. For example, PFBA and PFOA had a higher number of accumulated genes from 10-day compared to 1-day exposure (i.e., 10-day genes with BMCs = 288, and 550, respectively and 1-day BMCs = 113 and 215, respectively). PFSAs also exhibited a trend of more genes with lower BMCs with increasing chain-length; PFOS was the most bioactive affecting the largest number of genes prior to inducing cytotoxicity. PFOS is of concern to regulatory agencies due to its long historical use, environmental persistence, and long half-life of approximately 5.4 years in humans (Olsen *et al.*, 2007) and is one of the most prominent PFAS in human blood (Olsen *et al.*, 2017; Haines *et al.*, 2017). PFDS was more bioactive at low concentrations and was quite similar to PFHpS. In addition to increased transcriptional activity, the gene accumulation curves of these PFCAs and PFSAs shifted left, further indicating increased potency when comparing 1-day to 10-day exposures. Except for the sulfonamide (PFOSA), a PFOS precursor chemical and the 5:3 carboxylic acid, the majority of fluorotelomers did not elicit much transcriptional activity. PFOSA induced notable activity across both exposure times, with twice as many gene BMCs at 10-days relative to 1-day exposure (158 versus 318 genes with BMCs at 1 and 10-days, respectively). The 5:3 Acid also induced more transcriptional activity after a 10-day exposure (genes with BMCs = 288). Similar to PFCAs and PFSAs, such PFAA precursor gene accumulation curves also shifted left when exposure length was increased from 1 to 10-days.

### Applying thresholds of potential liver injury for prioritization and potency comparison

We explored the use of a liver injury threshold derived from a published approach (# of genes with BMCs > 105 genes) to compare PFAS potencies (Ramaiahgari *et al.* 2019). To ensure that all PFAS were included within the liver-injury potency ranking, the analysis included the first cytotoxic concentration, unlike the above BMC analysis that included only non-cytotoxic concentrations. This ensured that the PFAS that could not be modeled because of overt cytotoxicity at low concentrations at the 10-day time point (e.g., PFNA, PFDA, and PFUnA) would not be excluded from this analysis. A total of 16 PFAS surpassed this threshold at either the 1 or 10-day time points. Consistent with the results above, we also found that a greater number of PFAS exceeded this threshold after 10 days treatment (15 of 23 PFAS) compared to the 1-day treatment (11 of 23) (Fig. 4). For this purpose we used the (Ramaiahgari *et al.*, 2019).

**Figure 4.**
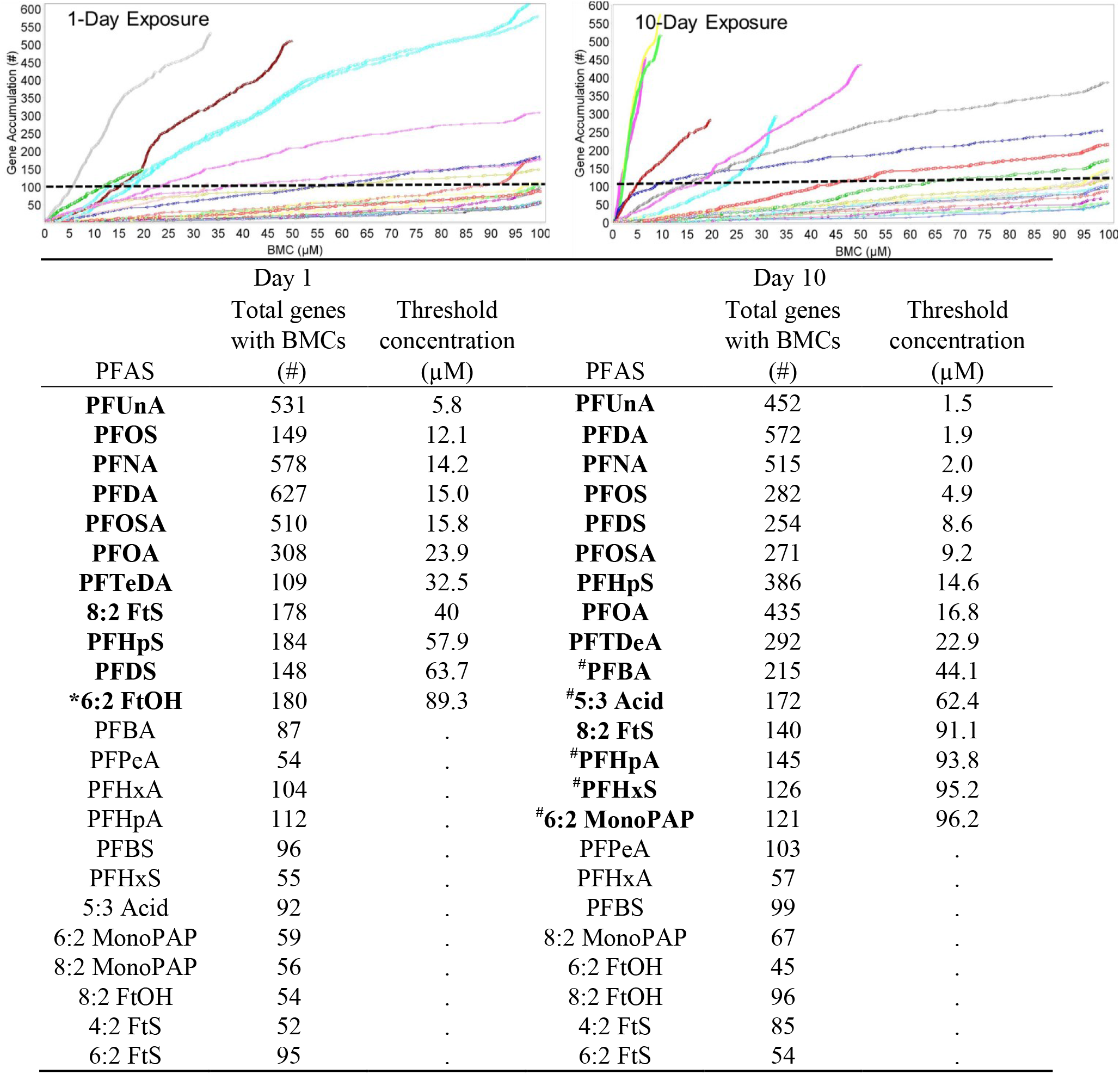
Threshold analysis and potency ranking using gene benchmark concentrations (BMCs) (adapted from Ramaiahgari et al. 2019) for PFAS after 1-day and 10-day exposure. Dashed line indicates a threshold of > 105 genes with BMCs proposed to differentiate chemicals that induce liver-injury (Bold). * PFAS surpassing the threshold from short-term (1-Day) but not long-term (10-day) exposure # PFAS i surpassing the threshold from long-term (10-Day) but not short-term (1-day) exposure

The concentration at which each PFAS surpassed the BMC liver-injury threshold was extrapolated from the gene accumulation plots (embedded table in Fig. 4), providing a surrogate measure of potency. The top four PFAS (PFUnA, PFOS, PFNA, and PFDA) became increasingly potent with time, surpassing the threshold at a lower concentration at 10-days (1.5, 4.9, 2.0, and 1.9 μM) compared to 1-day (5.8, 12.1, 14.2, and 15.5 μM) exposure, emphasizing the importance of our time series design. Based on this analysis, PFOS had a similar potency to PFUnA, which was consistent with the previous analysis (Fig. 3). When compared with the cytotoxicity analysis (Fig. 1), those most cytotoxic PFAS (i.e., inducing LDH leakage at lower concentrations), including PFNA, PFDA, PFUnA, PFOS and PFOSA, were the same PFAS that were considered to have the most potential to induce liver-injury based on the potency ranking (Fig. 4).

Ramaiahgari *et al.*, 2019 used a data-driven approach in establishing a threshold (# genes with BMC ≥ 105) for identification of liver injury inducing compounds in 2D HepaRG cultures. Although we are not able to predict ‘liver toxic’ chemicals as we did not have an equivalent training set for our liver spheroids, we applied this threshold approach instead to explore its utility in prioritizing and comparing potencies of the PFAS from our primary human liver cell spheroid data. Several PFAS surpassed the 105 gene BMC threshold across both timepoints, including PFCAs (PFOA, PFNA, PFDA, PFUnA) and PFSAs (PFHpS, PFOS, PFDS), as well as 8:2 FtS and PFOSA. Within these, we found that potency ranking based on the concentration at which the threshold was surpassed was consistent with the median BMC potency ranking. Thus, this approach allowed the qualitative separation of the most to the least potent PFAS *in vitro* for the purpose of prioritization and risk assessment.

We note that this threshold (# genes with BMC ≥ 105) could be increased to 150 or decreased to 75, resulting in a decrease or increase in the number of PFAS passing the thresholds, respectively, but without change to the potency ranking. Thus, application of this analysis approach to gene BMCs provides a robust methodology to rank PFAS potencies and prioritize chemicals for further research. This approach shows promise and could be more informative through the establishment of a liver toxic and non-toxic training set to further define hazard, as in Ramaiahgari *et al.* (2019).

### Deriving transcriptomic points of departure

The PFAS that were prioritized based on surpassing the 105 genes with BMCs threshold were subjected to further analysis as candidates for risk assessment by deriving a transcriptomic POD (Fig. 5). A conservative assessment using the BMC +/− BMCL and BMCU of the lower-bound 5^th^ percentile rounded to the nearest gene from the previous overall BMC analysis (Fig. 2) was used as the transcriptomic PODs for each PFAS depicted in Figure 5. This is consistent with previous approaches to deriving a molecular-based POD (Friedman *et al.*, 2019). Comparing 1- and 10-day exposures, a general trend of increasing potency was observed with decreased gene BMCs at 10 days, primarily with PFSAs. At the 1-day exposure time point, PFUnA, PFOS, and PFDS had the lowest mean BMCs with small confidence intervals, which was consistent with the overall median gene BMC approach. At the 10-day exposure time point, PFSAs showed a pattern of increasing potency with increasing chain-length. An exception to this is fluorotelomer 6:2 FtOH, on day 1, which was less potent, had wide confidence intervals, and did not meet the liver-toxic threshold on day 10. Overall, potency ranking was generally consistent for the 5^th^ percentile gene relative to the other approaches and provides a conservative BMC that could be used to inform a POD through *in vitro* to *in vivo* extrapolation when the required data for this application are produced.

**Figure 5.**
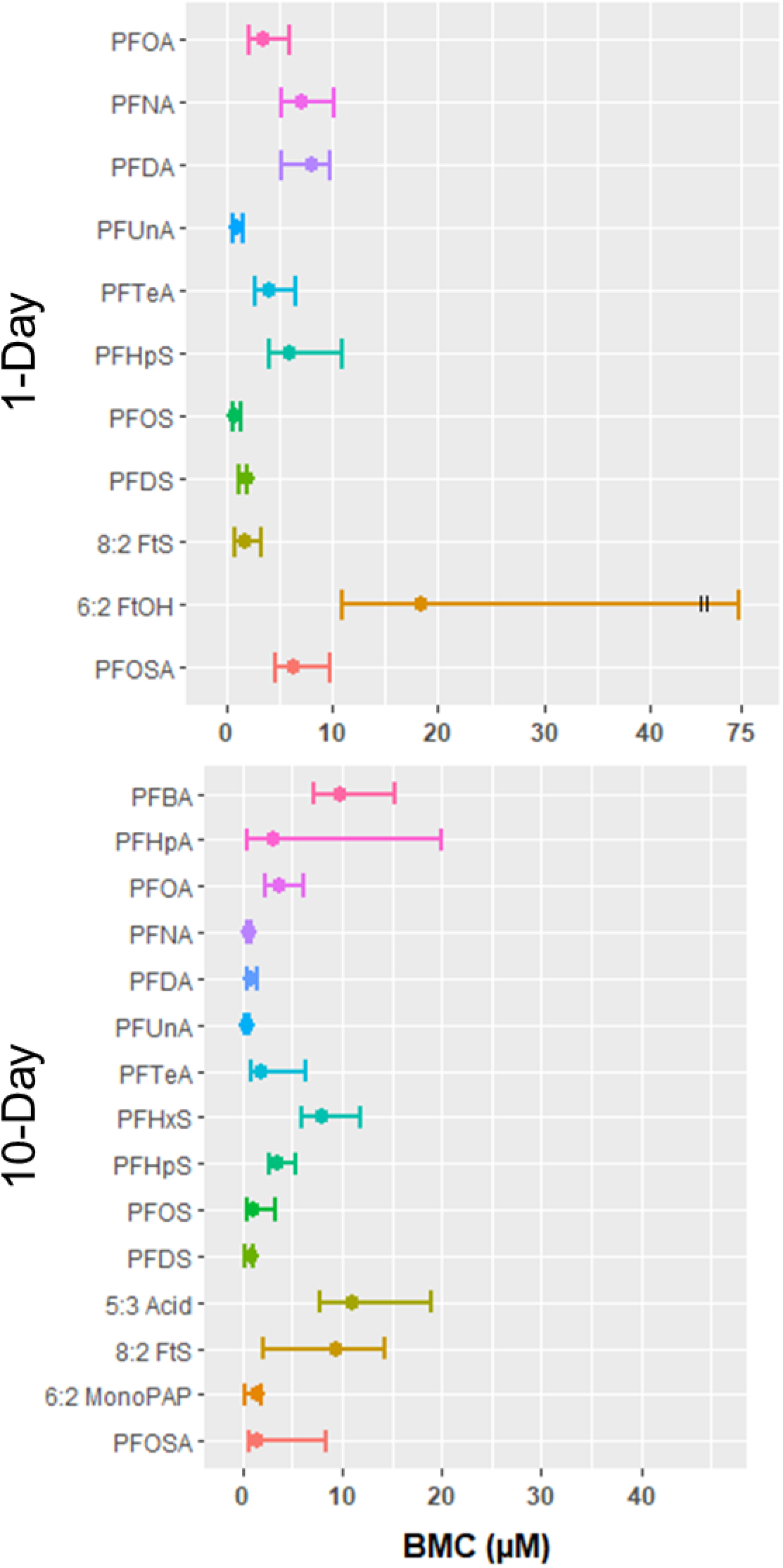
Range plots (median +/− 95 % confidence interval) of the lower bound 5^th^ percentile gene after 1-day, and 10-day PFAS exposure

## Discussion

This investigation extended our previous work (Rowan-Carroll *et al.* submitted) to include the broader class of PFAS. Here, the focus was to explore the magnitude of transcriptional changes as a measure of biological response to PFAS exposure. Thus, mode of action analyses (e.g., pathway analyses) were beyond the scope of this study. Our work demonstrates a robust high-content approach for toxicological assessment without the use of animal models, supporting the directive to reduce and eliminate the use of mammalian models in the future (e.g., US EPA directive (Wheeler, 2019)).

Using primary human liver cell spheroids as a model of toxicity, we found that longer-chain PFCAs, PFSAs, and PFOSA were more potent, and induced more transcriptional activity (more DEGs) and cytotoxicity than shorter chain. Moreover, these effects were more pronounced after longer exposures (10-days compared to 1-day). Decreased viability with increasing chain length has been reported previously in human liver HepG2 cells (Buhrke *et al.*, 2013; Ojo *et al.*, 2020), and also observed in human colon carcinoma cells exposed to PFAS carboxylic acids (Kleszczyński *et al.*, 2007). Observed trends in DEGs were also consistent with previous publications on PFOA and PFOS, which induce more prominent effects on gene expression following prolonged or chronic exposures (Rylander *et al.*, 2011; Fletcher *et al.*, 2013). Our findings suggest that structure (i.e., aliphatic chain length) plays a role in induced cytotoxicity from PFAS exposure in liver cell spheroids.

Recently, the National Toxicology Program (NTP) described modeling of toxicogenomic concentration-response data for use in chemical prioritization (Auerbach *et al.*, 2018). The goal of performing gene concentration-response studies was stated as “first and foremost to identify biological potency” as the names, descriptions, and establishment of phenotypic links with curated gene-sets might be misleading when interpreting toxicological effects (Auerbach *et al.*, 2018). Thus, this type of transcriptional concentration-response modeling is intended to be without reference to mode-of-action. Here, potency comparisons were first based on the median transcriptomic BMC, which we considered as similar to the AC50 (concentration at 50% of maximum activity). Although this approach is not conservative enough to be used to inform a transcriptomic POD, it is informative for chemical potency analysis of broad classes of environmental contaminants without attempting to predict specific hazards or mode of action.

BMC analyses supported observed results that PFCAs and PFSAs increase in potency and transcriptional activity with increasing carbon chain length. This trend of increased chemical potency with chain-length had been observed previously for PFCAs *in vitro* (Kleszczyński *et al.*, 2007), wherein specific PFAS accumulate within membrane bilayers with similar efficacy to cationic amphiphilic drugs that are known for their accumulative and lipid binding properties (Sanchez Garcia *et al.*, 2018). It was proposed that concentration and time-dependent changes in intracellular pH and acidification were the result of changes in membrane polarization based on aliphatic chain-length (Kleszczyński and Składanowski, 2009). Given the historical use of PFAS as a surfactant it is not surprising that prior studies have focused on PFAS distribution and accumulation, as well as resultant effects on specific properties (e.g., membrane polarization) of phospholipid bilayers (Hu *et al.*, 2003; Xie *et al.*, 2010). Such effects from PFAS membrane accumulation are consistent with short exposure (1-day) observations, but may not be sufficient to explain effects over longer exposures (10-days). However, organic anion transporters (e.g., OATPs, OAT2) are known to actively transport PFAS across the cell membrane into the cytosol (Zhao *et al.*, 2017) that may lead to accumulation of specific PFAS over time, exacerbated by the replenishing of PFAS-spiked media in spheroid cultures. Furthermore, PFOS is known to bind to human serum albumin, which is produced by hepatocytes (Chen and Guo, 2009; Zhang *et al.*, 2009). Although studies of the membrane kinetics and accumulation of PFAS were beyond the scope of the current investigation, chain-length dependent increases in potency and molecular activity from PFAS exposure could be used as a basis for chemical prioritization outside of an identified mechanism.

Overall, PFOS, PFUnA, and PFOSA were the most potent based on having the lowest transcriptional BMCs, and although PFNA and PFDA were less potent at 1-day, these chemicals were also some of the most potent PFAS at 10-days. PFNA, PFDA, and PFUnA also have longer elimination half-lives in humans when compared to PFOA (Han *et al.*, 2012; Zhang *et al.*, 2013) posing potential implications in humans from prolonged exposure. For example, whereas temporal trends of PFOA have been in historical decline, PFNA levels in the US population have changed very little from 2000 to 2015 (Olsen *et al.*, 2012, 2017), and these levels are also comparable to the Canadian population levels (monitored 2009 to 2011) (Haines *et al.*, 2017). Such prolonged exposure and accumulation of PFAS may be a contributing factor to PFAS potency when considering human health risk assessment. PFOS and PFUnA had identical gene BMC accumulation plots in our study on day 1. Although PFUnA is detected in Canadian blood plasma, there are fewer data on the toxicological implications of its exposure relative to PFOS. In repeat dose-response studies in rats, the NOAEL was 0.1 mg/kg-bw/day for PFUnA (Takahashi *et al.*, 2014) compared to 0.02 mg/kg-bw/day for PFOS (Butenhoff *et al.*, 2012). Our results indicate that PFUnA may be equipotent to PFOS and other well-known PFAS in human liver cells based on transcriptional perturbations.

PFOSA, an intermediate metabolite of longer-chain PFOS precursors (e.g., N-ethylperfluorooctane sulfonamide, NEtFOSA), had similar potency and transcriptomic activity to PFOS. Prior animal studies have demonstrated the capacity of PFOSA to be metabolized into PFOS (Chen *et al.*, 2015; Ross *et al.*, 2012), and a significant portion of this metabolism takes place in the liver (Chen *et al.*, 2015). Such studies suggest precursor compounds may lead to indirect PFAS human exposure (Gebbink *et al.*, 2015). Our findings indicate that PFOSA elicits changes in gene expression with the same potency and transcriptional activity as PFOS. Given the metabolic proficiency of 3D spheroid models, it is reasonable to suggest that PFOSA exposures may yield fractional levels of PFOS that contribute to these similarities.

The approach proposed by the NTP applies genomic concentration-response modeling to identify the most sensitive median gene-set BMC, referred to as the biological effect POD (i.e., the lowest concentration that induces biological effects) in short-term animal studies (Auerbach *et al.*, 2018). Thomas *et al.*, (2013) demonstrated that transcriptional changes from gene-sets such as the lowest pathway BMC in rodents is highly correlated with apical endpoints for numerous chemicals when considering both cancer and non-cancer endpoints. Furthermore, Farmahin *et al.*, (2017) showed that the method of selecting a subset of genes to derive a transcriptional POD is generally conservative and that a variety of approaches may be suitable for risk assessment. There are several potential issues associated with the use of gene-set BMCs; e.g., differences in the pathway databases used for the gene sets, the fact that genes can operate in multiple pathways, and subjectivity in setting thresholds for gene-set BMC derivation. Thus, we explored the use of the lower-bound 5^th^ percentile of all genes with BMCs rounded to the nearest gene for each PFAS at each exposure considered to determine a transcriptional POD. Interestingly, although these 5^th^ percentile BMCs are more conservative than the BMC medians (as is desired for a POD used in priority setting or screening), they provide a comparable *in vitro* potency rank order, reinforcing results from the median BMC approach, with lower PODs for longer chain lengths. In addition, one problem we identified with gene set BMCs was the very large confidence intervals for the lowest pathways because these tended to be based on very few genes (data not shown), which impaired the ability to observe significant differences between the PFAS. The 5^th^ percentile BMC, which is consistent with the approach used in the ToxCast program, resolved this issue. Overall, the PFAS with PODs equivalent to (overlapping confidence intervals), or below, PFOS and PFOA were: PFUnA, PFDS, 8:2 Fts, and PFNA, PFDA, PFUnA, PFDS, 6:2 monoPAP, for 1-day and 10-day exposures, respectively.

The next step would be to derive the appropriate data to perform *in vitro* to *in vivo* extrapolation modeling and determine a human AED. Within Rowan-Carroll *et al.* (submitted), we used this approach to calculate bioactivity exposure ratios (BERs) for PFOS and PFOA and found that they were within range (e.g., 2-orders of magnitude) of daily human exposure levels. These BER values were considered low enough to indicate higher risk to human health and warrant attention for future risk assessment. Transcriptomic BER analysis represents a novel approach for use in current risk assessment practices providing an approach for analysis of data-poor chemicals that is powerful within the context of a tiered framework approach for evaluating risk from exposure to environmental chemicals (Gannon *et al.*, 2019). Moreover, an increased recognition of the contributions of bioaccumulation in these spheroid models with PFAS exposure enables an improved understanding of the intrinsically interdependent effects of bioaccumulation, metabolism/transport, and biological responses for refined translation to humans through *in vitro* to *in vivo* extrapolation.

## Conclusion

Our work demonstrates how BMC modeling can be used for prioritization, potency assessment, and identification of an *in vitro* transcriptomic POD. We found a number of PFAS that cause cytotoxicity in human liver cell spheroids and transcriptional changes at similar or concentrations to PFOS and PFOA, suggesting these chemicals are harmful to human liver. A clear relationship between chain length and extent of transcriptional alterations, and potency, emerged that can be used to inform read-across. Future work will explore mode of action in relation to functional groups to enhance grouping for read-across.

## Supporting information

Supplemental Information for Reardon et al.

## Acknowledgements

Thanks to Dr. Barbara Wetmore for providing the PFAS and accompanying QA/QC data for the stocks procured under contract to the EPA. Thanks to Dr. Scott Auerbach of the Toxicoinformatics Group within the National Toxicology Program (NIEHS) for reviewing anf providing valuable feedback on the manuscript.

