## Supplemental Information for Reardon et al. for "High-throughput transcriptomics and benchmark concentration modeling for potency ranking of per- and polyfluoroalkyl substances (PFAS) in exposed human liver cell spheroids"

**Supplementary Information**

TempOSeq Data – Data generation, normalization, and plate-effect correction

***DMB data generation***

Low count probes, i.e. less than a median of 5 counts within an exposure group for a probe, were eliminated from the dataset. Groups having a higher percentage of DMSO {0.17, 0.3} were normalized on the cpm scale to the most common group {0.1} by applying the following for each probe;

*Example calculations:*

For DMSO0.3

newProbeLog2CPMvalue = log2(probeCPMvalue) - [log2(mean(DMSO0.3)) – log2(mean (DMSO0.1)]

For DMSO0.17

newProbeLog2CPMvalue = log2(probeCPMvalue) - [log2(mean(DMSO0.17)) – log2(mean (DMSO0.1)]

If a plate correction was made:

Data were 2x, then

newProbeLog2CPMvalue = log2(probeCPMvalue) - [log2(mean(DMSO0.1ForPlateX)) – log2(mean(DMSO0.1ForAllPlates))]

***Plate-effect correction***

Low count probes, i.e. less than a median of 5 counts within an exposure group for a probe, were flagged as “absent” from the dataset. The Differential Gene Expression (DGE) analysis conducted with the default parameters of DESeq2 v1.24 with respective control and exposure groups. Probes reaching the threshold of an adjusted p-value <0.05 and an absolute fold change > 1.5 were flagged (probes passing filters; PPF) and were retained for further analyses.

Treatment of genes fitting “flagged” Hill models

Hill models were flagged if the “k” parameter is less than 1/3 the lowest positive treatment concentration (e.g., 0.2 µM). In this case the next best model with a goodness-of-fit p-value < 0.05 is selected. However, if no model is fit (have a p-value of > 0.05) then genes that fit Hill models (excluding flagged models) were included, and the lowest benchmark dose lower confidence limit (BMC_L_) is considered and divided by 2 for the final analysis.

Idenitfying liver injury from BMC gene accumulation plots from established thresholds of liver injury

BMC accumulation plots have also been used for hazard identification of potentially liver-toxic compounds. Ramaiahgari et al. (2019) described a novel approach deriving a liver injury threshold to distinguish liver-toxic from non-toxic compounds. They produced gene BMCs from HepaRG cells exposed to six low-risk, non-liver-cytotoxic chemicals (caffeine, sucrose, KCl, levofloxacin, phenobarbital, and aspirin) as a measure of transcriptional activity. They used a cutoff of 3 SDs above the mean number of gene BMCs from these non-hepatotoxic chemicals as the threshold. Chemicals surpassing this threshold (# genes with BMCs ≥ 105) were expected to induce liver toxicity, and the concentration at which BMC accumulation exceeded this bootstrapped threshold was proposed as a quantitative estimate for liver injury. Thus, they were able to distinguish established hepatotoxic chemicals from compounds rarely associated with hepatotoxicity (Ramaiahgari et al., 2019).

**Table S1**. PFAS induced cytotoxicity in liver spheroids from 1 and 10-Day exposures

|  | PFAS | Highest non-cytotoxic concentration (µM) | Concentrations removed from DEG and BMC analyses due to cytotoxicity (µM) |
| --- | --- | --- | --- |
| 1-Day Exposure | PFDA | 50 | 100 |
|  | PFUnA | 13 | 34, 66 |
|  | PFOS | 20 | 50, 100 |
|  | PFOSA | 20 | 50, 100 |
| 10-Day Exposure | PFOA | 50 | 100 |
|  | PFNA | 2 | 10, 20, 50, 100 |
|  | PFDA | 2 | 10, 20, 50, 100 |
|  | PFUnA | 1.3 | 6.5, 13, 34, 66 |
|  | PFTeA | 17 | 33 |
|  | PFOS | 20 | 50, 100 |
|  | PFOSA | 20 | 50, 100 |
|  | 8:2 FtS | 50 | 100 |

**Table S2.** Differentially expressed genes and concentration-responses of human liver spheroids after 1 and 10-day exposure to PFAS

|  | **Day** | **Concentration (µM)** | | | | | | **Trend** |
| --- | --- | --- | --- | --- | --- | --- | --- | --- |
|  |  | **0.2** | **2** | **10** | **20** | **50** | **100** |  |
| PFBA | 1 | 5 | 42 | 1 | 10 | 15 | 58 |  |
|  | 10 | 41 | 49 | 17 | 19 | 132 | 199 |  |
| PFPeA | 1 | 0 | 1 | 2 | 2 | 0 | 0 |  |
|  | 10 | 34 | 40 | 22 | 40 | 26 | 24 |  |
| PFHxA | 1 | 119 | 43 | 22 | 60 | 101 | 24 |  |
|  | 10 | 123 | 167 | 139 | 61 | 44 | 62 |  |
| PFHpA | 1 | 0 | 0 | 1 | 4 | 14 | 51 |  |
|  | 10 | 55 | 10 | 19 | 25 | 57 | 91 |  |
| PFOA | 1 | 0 | 11 | 72 | 71 | 229 | 491 |  |
|  | 10 | 2 | 11 | 85 | 106 | 625 | -* |  |
| PFNA | 1 | 3 | 0 | 37 | 167 | 236 | 785 |  |
|  | 10 | 165 | 36 | -* | -* | -* | -* |  |
| PFDA | 1 | 0 | 4 | 1 | 70 | 364 | -* |  |
|  | 10 | 29 | 34 | -* | -* | -* | -* |  |
| PFUnA^A^ | 1 | 40 | 20 | 119 | 227 | -* | -* |  |
|  | 10 | 0 | 0 | -* | -* | -* | -* |  |
| PFTeDA^B^ | 1 | 46 | 9 | 128 | 55 | 69 | 43 |  |
|  | 10 | 42 | 0 | 67 | 47 | 67 | -* |  |
| PFBS | 1 | 0 | 34 | 7 | 0 | 50 | 72 |  |
|  | 10 | 0 | 2 | 0 | 7 | 55 | 79 |  |
| PFHxS | 1 | 47 | 0 | 0 | 12 | 11 | 16 |  |
|  | 10 | 60 | 22 | 8 | 1 | 0 | 2 |  |
| PFHpS | 1 | 0 | 1 | 14 | 26 | 61 | 225 |  |
|  | 10 | 80 | 10 | 93 | 132 | 262 | 467 |  |
| PFOS | 1 | 1 | 50 | 171 | 295 | -* | -* |  |
|  | 10 | 4 | 64 | 175 | 498 | -* | -* |  |
| PFDS | 1 | 0 | 17 | 49 | 75 | 177 | 190 |  |
|  | 10 | 0 | 42 | 140 | 181 | 241 | 237 |  |
| 5:3 Acid | 1 | 0 | 3 | 0 | 9 | 8 | 23 |  |
|  | 10 | 15 | 3 | 6 | 6 | 20 | 131 |  |
| 6:2 MonoPAP | 1 | 0 | 0 | 7 | 190 | 6 | 7 |  |
|  | 10 | 5 | 17 | 15 | 27 | 30 | 20 |  |
| 8:2 MonoPAP | 1 | 1 | 0 | 0 | 0 | 1 | 3 |  |
|  | 10 | 6 | 0 | 0 | 0 | 0 | 11 |  |
| 6:2 FtOH | 1 | 12 | 55 | 43 | 21 | 67 | 32 |  |
|  | 10 | 93 | 41 | 107 | 22 | 10 | 0 |  |
| 8:2 FtOH | 1 | 0 | 7 | 0 | 9 | 0 | 0 |  |
|  | 10 | 37 | 20 | 13 | 31 | 20 | 72 |  |
| 4:2 FTS | 1 | - | 10 | 19 | 10 | 5 | 1 |  |
|  | 10 | 47 | 40 | 71 | 31 | 33 | 43 |  |
| 6:2 FtS | 1 | 3 | 5 | 15 | 6 | 69 | 7 |  |
|  | 10 | 34 | 40 | 12 | 10 | 16 | 51 |  |
| 8:2 FtS | 1 | 4 | 29 | 56 | 127 | 212 | 237 |  |
|  | 10 | 81 | 88 | 136 | 172 | 234 | -* |  |
| PFOSA | 1 | 10 | 30 | 18 | 141 | -* | -* |  |
|  | 10 | 38 | 102 | 93 | 488 | -* | -* |  |

|  | PFBA |
| --- | --- |
| 0.2 | 0.69897 |
| 2 | 1.62325 |
| 10 | 0 |
| 20 | 1 |
| 50 | 1.17609 |
| 100 | 1.76343 |

|  | PFBA |
| --- | --- |
| 0.2 | 1.61278 |
| 2 | 1.6902 |
| 10 | 1.23045 |
| 20 | 1.27875 |
| 50 | 2.12057 |
| 100 | 2.29885 |

|  | PFPeA |
| --- | --- |
| 0.2 | 1.716 |
| 2 | 0 |
| 10 | 0.30103 |
| 20 | 0.30103 |
| 50 | 0 |
| 100 | 0 |

|  | PFPeA |
| --- | --- |
| 0.2 | 1.53148 |
| 2 | 1.60206 |
| 10 | 1.34242 |
| 20 | 1.60206 |
| 50 | 1.41497 |
| 100 | 1.38021 |

|  | PFHxA |
| --- | --- |
| 0.2 | 2.07555 |
| 2 | 1.63347 |
| 10 | 1.34242 |
| 20 | 1.77815 |
| 50 | 2.00432 |
| 100 | 1.38021 |

|  | PFHxA |
| --- | --- |
| 0.2 | 2.08991 |
| 2 | 2.22272 |
| 10 | 2.14301 |
| 20 | 1.78533 |
| 50 | 1.64345 |
| 100 | 1.79239 |

|  | PFHpA |
| --- | --- |
| 0.2 | 1.20412 |
| 2 | 0 |
| 10 | 0.477121 |
| 20 | 0 |
| 50 | 1.17609 |
| 100 | 1.85733 |

|  | PFHpA |
| --- | --- |
| 0.2 | 1.74036 |
| 2 | 1 |
| 10 | 1.27875 |
| 20 | 1.39794 |
| 50 | 1.75587 |
| 100 | 1.95904 |

|  | PFOA |
| --- | --- |
| 0.2 | 1.43136 |
| 2 | 1.04139 |
| 10 | 1.85733 |
| 20 | 1.85126 |
| 50 | 2.35984 |
| 100 | 2.69108 |

|  | PFOA |
| --- | --- |
| 0.2 | 0.30103 |
| 2 | 1.04139 |
| 10 | 1.92942 |
| 20 | 2.02531 |
| 50 | 2.79588 |

|  | PFNA |
| --- | --- |
| 0.2 | 0.477121 |
| 2 | 0 |
| 10 | 1.5682 |
| 20 | 2.22272 |
| 50 | 2.37291 |
| 100 | 2.89487 |

|  | PFNA |
| --- | --- |
| 0.2 | 2.21748 |
| 2 | 1.5563 |

|  | PFDA |
| --- | --- |
| 0.2 | 0 |
| 2 | 0.60206 |
| 10 | 0 |
| 20 | 1.79239 |
| 50 | 2.54777 |

|  | PFDA |
| --- | --- |
| 0.2 | 1.4624 |
| 2 | 1.53148 |

|  | PFUnA |
| --- | --- |
| 0.13 | 1.60206 |
| 1.3 | 1.30103 |
| 6.5 | 2.07555 |
| 13 | 2.35603 |

|  | PFUnA |
| --- | --- |
| 0.13 | 0 |
| 1.3 | 0 |

|  | PFTeDA |
| --- | --- |
| 0.06 | 1.66276 |
| 0.67 | 0.954243 |
| 3.35 | 2.10721 |
| 6.7 | 1.74036 |
| 17 | 1.83885 |
| 33 | 1.63347 |

|  | PFTeDA |
| --- | --- |
| 0.06 | 1.62325 |
| 0.67 | 0 |
| 3.35 | 1.82607 |
| 6.7 | 1.6721 |
| 17 | 1.82607 |

|  | PFBS |
| --- | --- |
| 0.2 | 0 |
| 2 | 1.53148 |
| 10 | 0.845098 |
| 20 | 0 |
| 50 | 1.69897 |
| 100 | 1.85733 |

|  | PFBS |
| --- | --- |
| 0.2 | 0 |
| 2 | 0.30103 |
| 10 | 0 |
| 20 | 0.845098 |
| 50 | 1.74036 |
| 100 | 1.89763 |

|  | PFHxS |
| --- | --- |
| 0.2 | 1.6721 |
| 2 | 0 |
| 10 | 0 |
| 20 | 1.07918 |
| 50 | 1.04139 |
| 100 | 1.20412 |

|  | PFHxS |
| --- | --- |
| 0.2 | 1.77815 |
| 2 | 1.34242 |
| 10 | 0.90309 |
| 20 | 0 |
| 50 | 0 |
| 100 | 0.30103 |

|  | PFHpS |
| --- | --- |
| 0.2 | 0 |
| 2 | 0 |
| 10 | 1.14613 |
| 20 | 1.41497 |
| 50 | 1.78533 |
| 100 | 2.35218 |

|  | PFHpS |
| --- | --- |
| 0.2 | 1.90309 |
| 2 | 1 |
| 10 | 1.96848 |
| 20 | 2.12057 |
| 50 | 2.4183 |
| 100 | 2.66932 |

|  | PFOS |
| --- | --- |
| 0.2 | 0 |
| 2 | 1.69897 |
| 10 | 2.233 |
| 20 | 2.46982 |

|  | PFOS |
| --- | --- |
| 0.2 | 0.60206 |
| 2 | 1.80618 |
| 10 | 2.24304 |
| 20 | 2.69723 |

|  | PFDS |
| --- | --- |
| 0.2 | 0 |
| 2 | 1.23045 |
| 10 | 1.6902 |
| 20 | 1.87506 |
| 50 | 2.24797 |
| 100 | 2.27875 |

|  | PFDS |
| --- | --- |
| 0.2 | 0 |
| 2 | 1.62325 |
| 10 | 2.14613 |
| 20 | 2.25768 |
| 50 | 2.38202 |
| 100 | 2.37475 |

|  | c53Acid |
| --- | --- |
| 0.2 | 0 |
| 2 | 0.477121 |
| 10 | 0 |
| 20 | 0.954243 |
| 50 | 0.90309 |
| 100 | 1.36173 |

|  | c53Acid |
| --- | --- |
| 0.2 | 1.17609 |
| 2 | 0.477121 |
| 10 | 0.778151 |
| 20 | 0.778151 |
| 50 | 1.30103 |
| 100 | 2.11727 |

|  | C62monoPAP |
| --- | --- |
| 0.2 | 0 |
| 2 | 0 |
| 10 | 1.44716 |
| 20 | 0 |
| 50 | 0.845098 |
| 100 | 1.04139 |

|  | C62monoPAP |
| --- | --- |
| 0.2 | 0.69897 |
| 2 | 1.23045 |
| 10 | 1.17609 |
| 20 | 1.43136 |
| 50 | 1.47712 |
| 100 | 1.30103 |

|  | C82MonoPAP |
| --- | --- |
| 0.2 | 0 |
| 2 | 0 |
| 10 | 0 |
| 20 | 0 |
| 50 | 0 |
| 100 | 0.477121 |

|  | C82MonoPAP |
| --- | --- |
| 0.2 | 0.778151 |
| 2 | 0 |
| 10 | 0 |
| 20 | 0 |
| 50 | 0 |
| 100 | 1.04139 |

|  | FtOH62 |
| --- | --- |
| 0.2 | 1.07918 |
| 2 | 1.74036 |
| 10 | 1.63347 |
| 20 | 1.32222 |
| 50 | 1.82607 |
| 100 | 1.50515 |

|  | FtOH62 |
| --- | --- |
| 0.2 | 1.96848 |
| 2 | 1.61278 |
| 10 | 2.02938 |
| 20 | 1.34242 |
| 50 | 1 |
| 100 | 0 |

|  | FtOH82 |
| --- | --- |
| 0.2 | 0 |
| 2 | 0.845098 |
| 10 | 0 |
| 20 | 0.954243 |
| 50 | 0 |
| 100 | 0 |

|  | FtOH82 |
| --- | --- |
| 0.2 | 1.5682 |
| 2 | 1.30103 |
| 10 | 1.11394 |
| 20 | 1.49136 |
| 50 | 1.30103 |
| 100 | 1.85733 |

|  | FTS42 |
| --- | --- |
| 0.2 | 0 |
| 2 | 1 |
| 10 | 1.27875 |
| 20 | 1 |
| 50 | 0.69897 |
| 100 | 0 |

|  | FTS42 |
| --- | --- |
| 0.2 | 1.6721 |
| 2 | 1.60206 |
| 10 | 1.85126 |
| 20 | 1.49136 |
| 50 | 1.51851 |
| 100 | 1.63347 |

|  | FtS62 |
| --- | --- |
| 0.2 | 0.477121 |
| 2 | 0.69897 |
| 10 | 1.17609 |
| 20 | 0.778151 |
| 50 | 1.83885 |
| 100 | 0.845098 |

|  | FtS62 |
| --- | --- |
| 0.2 | 1.53148 |
| 2 | 1.60206 |
| 10 | 1.07918 |
| 20 | 1 |
| 50 | 1.20412 |
| 100 | 1.70757 |

|  | FtS82 |
| --- | --- |
| 0.2 | 0.60206 |
| 2 | 1.4624 |
| 10 | 1.74819 |
| 20 | 2.1038 |
| 50 | 2.32634 |
| 100 | 2.37475 |

|  | FtS82 |
| --- | --- |
| 0.2 | 1.90849 |
| 2 | 1.94448 |
| 10 | 2.13354 |
| 20 | 2.23553 |
| 50 | 2.36922 |

|  | *PFOSA |
| --- | --- |
| 0.2 | 1 |
| 2 | 1.47712 |
| 10 | 1.25527 |
| 20 | 2.14922 |

|  | *PFOSA |
| --- | --- |
| 0.2 | 1.57978 |
| 2 | 2.0086 |
| 10 | 1.96848 |
| 20 | 2.68842 |

* Overtly cytotoxic concentrations eliminated from analysis,
 ^A^ Concentration range of PFUnA (0.13, 1.3, 6.5, 13, 34, 66 µM),
 ^B^ Concentration range of PFTeDA (0.06, 0.67, 3.35, 6.7, 17, 33 µM)

**Table S3**. Benchmark concentration (median BMC ± 95 % CI) from 1 and 10-Day exposure to all measured PFAS

|  | **PFAS** | **1-Day**  median BMC  (Lower, Upper 95 % CI) | **10-Day**  median BMC  (Lower, Upper 95 % CI) |
| --- | --- | --- | --- |
| PFCAs | PFBA | 56.5 (54.2, 64.3) | 40.4 (36.0, 43.7) |
|  | PFPeA | 74.8 (71.1, 85.5) | 59.3 (53.4, 68.4) |
|  | PFHxA | 64.3 (58.4, 77.8) | 66.8 (41.1, 75.1) |
|  | PFHpA | 50.8 (48.5, 58.6) | 67.7 (56.8, 74.1) |
|  | PFOA | 38.9 (34.8, 43.2) | 31.2 (28.9, 33.9) |
|  | PFNA | 42.5 (40.7, 45.6) | 1.1 (1.0, 1.2) |
|  | PFDA | 23.1 (21.9, 24.5) | 1.2 (1.0, 1.3) |
|  | PFUnA | 7.0 (6.3, 7.6) | 1.0 (0.8, 1.3) |
|  | PFTeA | 19.9 (18.4, 23.7) | 14.3 (12.5, 15.7) |
| PFSAs | PFBS | 54.9 (51.8, 61.7) | 46.2 (42.3, 50.3) |
|  | PFHxS | 55.3 (40.8, 66.6) | 60.5 (55.7, 67.6) |
|  | PFHpS | 49.4 (41.9, 56.3) | 39.6 (36.0, 43.8) |
|  | PFOS | 7.3 (6.5, 8.3) | 8.1 (7.0, 9.0) |
|  | PFDS | 18.8 (15.2, 24.8) | 12.0, 10.3, 16.4) |
| Precursors | C5:3 Acid | 62.0 (53.3, 68.7) | 48.9 (45.6, 53.6) |
|  | 6:2 monoPAP | 60.8 (52.3, 68.3) | 50.0 (43.5, 56.7) |
|  | 8:2 monoPAP | 80.9 (71.5, 86.2) | 63.7 (56.2, 71.0) |
|  | 6:2 FtOH | 84.7 (77.3, 87.8) | 62.6 (49.4, 85.0) |
|  | 8:2 FtOH | 69.3 (63.3, 74.9) | 58.7 (53.2, 70.8) |
|  | 4:2 FtS | 61.8 (52.4, 72.8) | 64.5 (55.0, 71.7) |
|  | 6:2 FtS | 68.4 (48.7, 82.1) | 68.8 (62.7, 80.5) |
|  | 8:2 FtS | 35.5 (29.4, 41.4) | 37.8 (32.2, 43.5) |
|  | PFOSA | 13.4 (12.8, 14.3) | 11.5 (10.3, 12.7) |

**Figure S1**. Log-transformed gene accumulation plots of BMC responses in human liver spheroids following exposure to A) PFCAs B) PFSAs or C) fluorotelomers/sulfonamides exposed from 1-day (BMDExpress 2.3).

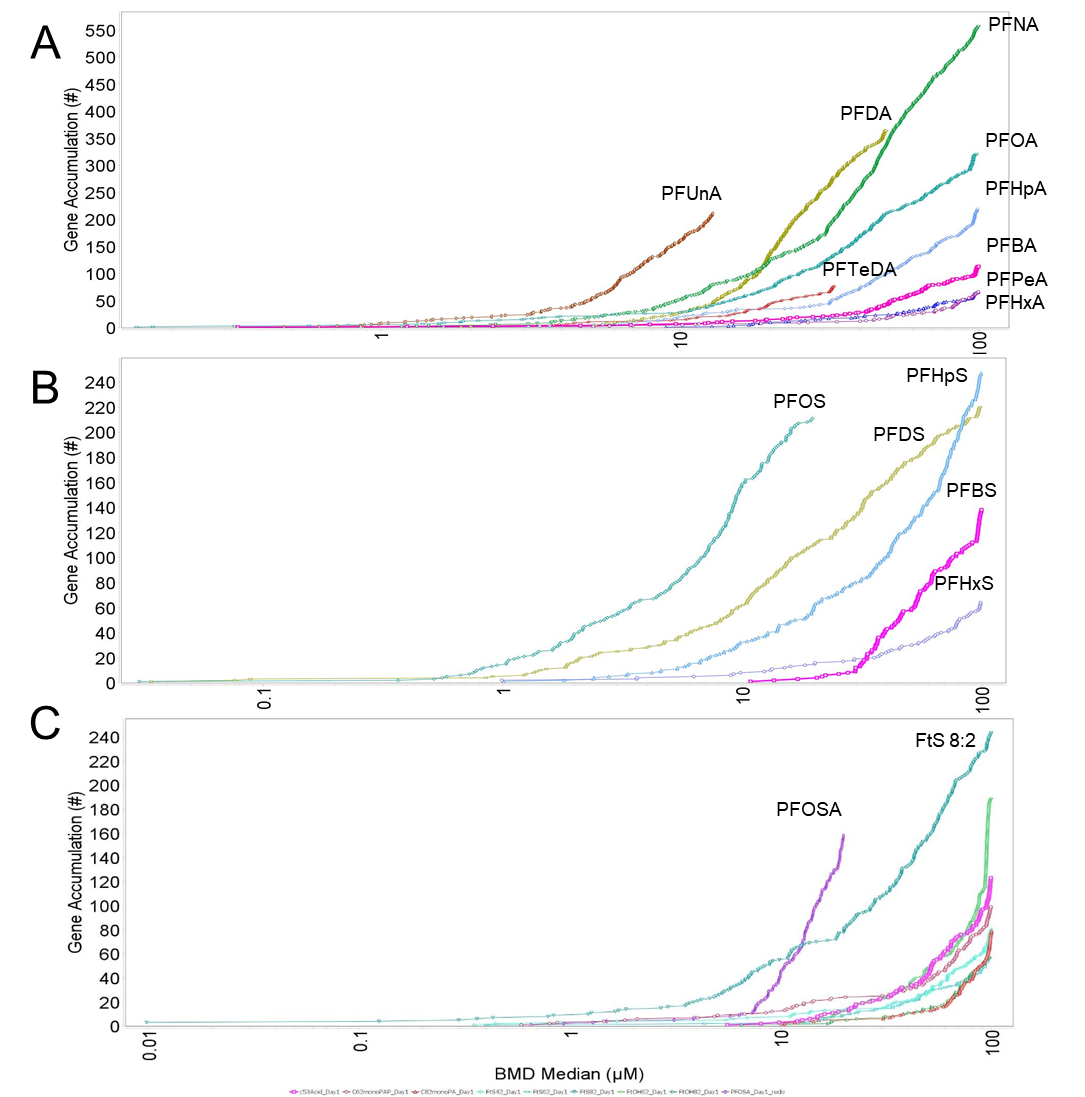

**Figure S2**. Log-transformed gene accumulation plots of BMC responses in human liver spheroids following exposure to A) PFCAs B) PFSAs or C) fluorotelomers/sulfonamides exposed from 10-day (BMDExpress 2.3).

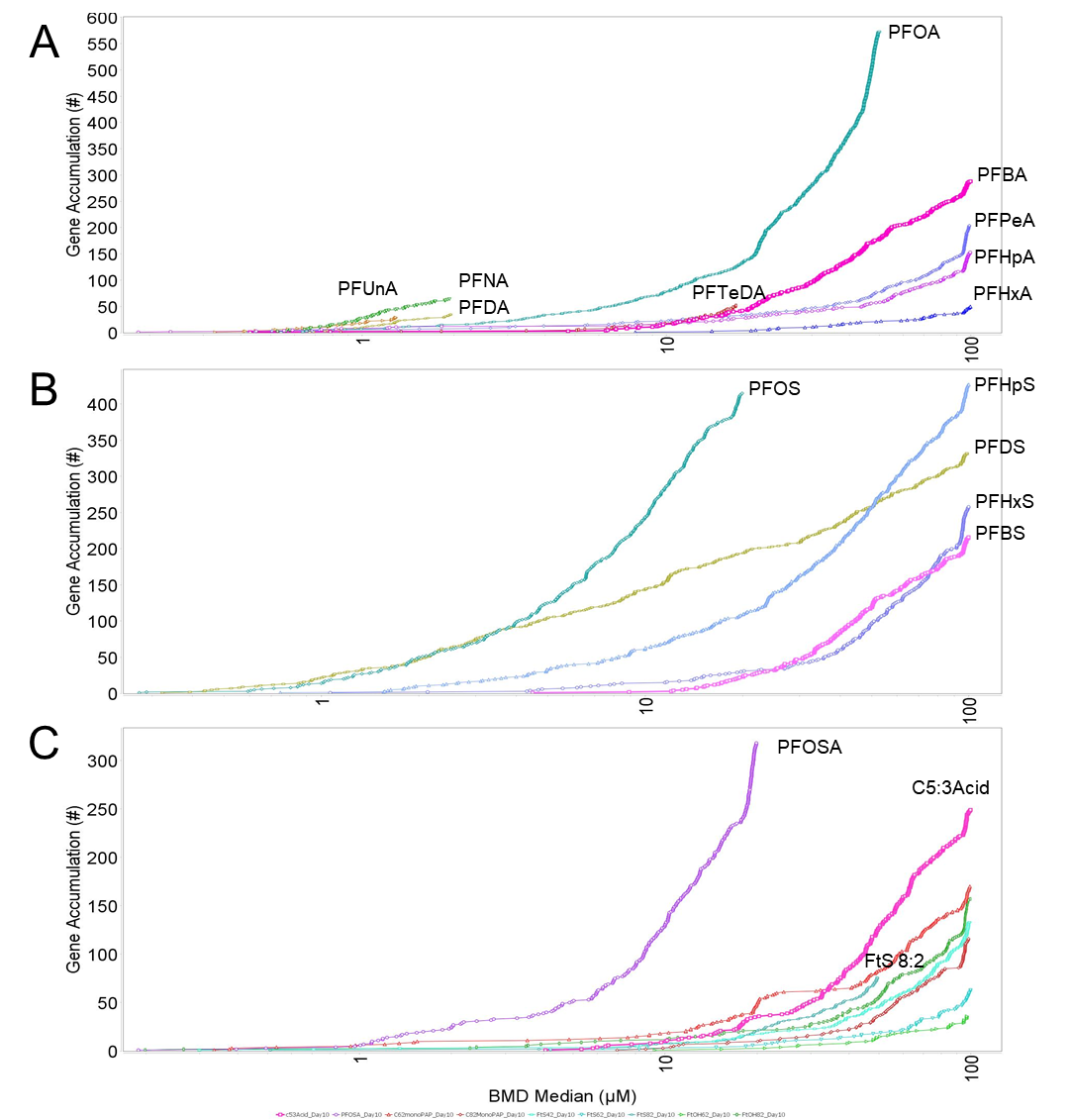
